# immunoPETE: A DNA-based integrated B-cell and T-cell receptor profiling platform

**DOI:** 10.64898/2026.03.17.712532

**Authors:** Hahn Zhao, Hamid Mirebrahim, Dilduz Telman, Richard Dannebaum, Sylvie McNamara, Ehsan Tabari, Hai Lin, Florian Rubelt, Jan Berka, Khai Luong, Magdalene Joseph, Richard Bryan, Douglas Ward, Adrian Hayday, Sowmi Utiramerur, Dinesh Kumar, Hosseinali Asgharian

**Affiliations:** Hoffmann-La Roche Limited, 7070 Mississauga Rd, Mississauga, ON L5N 5M8, Canada; CSI Roche Diagnostics Solutions, 4300 Hacienda Dr, Pleasanton, CA 94588; Roche Diagnostics Solutions, 4300 Hacienda Dr, Pleasanton, CA 94588; Department of Immunobiology, King’s College London, London, UK; Department of Cancer and Genomic Sciences, University of Birmingham, Birmingham, UK

**Author notes:** Equal Contribution (Hahn Zhao, Hamid Mirebrahim).

## Abstract

The vast diversity of B and T cell receptors generated through the recombination of Variable (V), Diversity (D), and Joining (J) gene segments plays a critical role in adaptive immunity. Profiling immune repertoires at the DNA level provides a robust and stable approach to capture the clonal composition of these receptors. immunoPETE is an assay designed to target recombined human T-cell Receptor Beta (TRB), T-cell Receptor Delta (TRD), and Immunoglobulin Heavy (IGH) chain genes directly from genomic DNA. Simultaneous profiling of B and T cell receptor chains in a single reaction provides internally normalized clone counts and facilitates the study of B-T cell interactions. Full-length amplicon consensus sequences representative of original template DNA molecules are accurately reconstructed using Unique Molecular Identifiers (UMIs). An in-house pipeline compiles VDJ rearrangements from the Complementarity-Determining Region 3 (CDR3) of TRB, TRD and IGH chains into comprehensive readouts at cell-level resolution. In this study, we describe the immunoPETE end-to-end workflow, followed by a comprehensive benchmarking of its performance in adaptive immune profiling. Where applicable, we used both natural and contrived samples and characterized the assay’s accuracy, linearity, and reproducibility across several metrics: retrieving CDR3 sequences, determining B and T cell ratios, total cell count, yield, fraction of functional rearrangements, clonal diversity, composition of dominant clones, pairwise similarity, and V/J gene usage frequencies. Furthermore, we assessed its quantitative limits concerning the total number of lymphocytes and the detection of rare clones. As an example of its applications, we show that adding immune biomarkers extracted from immunoPETE data to clinical factors improves prediction of progression-free survival in a cohort of non-muscle invasive bladder cancer (NMIBC) patients. Finally, we discuss the broad applications of immunoPETE in the study of aging, cancers, infections, and autoimmune disorders with reference to select published studies.

## 1. Introduction

The adaptive immune system relies on a sophisticated division of labor between B and T lymphocytes to provide highly specific, long-lasting protection. The precision of these mechanisms is driven by highly specific antigen receptors—T cell receptors (TCRs) and B cell receptors (BCRs)—which ensure targeted and efficient responses to a vast array of antigens. B cells serve as the humoral arm of immune defense, identifying extracellular pathogens and differentiating into plasma cells to secrete antibodies that neutralize threats. In contrast, T cells govern cell-mediated immunity through specialized subsets. Helper T cells (CD4^+^) act as the “command center,” secreting cytokines to coordinate the response of other immune cells, while Cytotoxic T cells (CD8^+^) are the primary effectors, directly seeking and attacking virally infected or cancerous cells by inducing apoptosis (Janeway, Travers, Walport, & Shlomchik, 2001a). Beyond these traditional roles, the system includes cells that bridge the gap between innate and adaptive responses: Gamma delta (γδ) T cells possess TCRs with limited diversity and often reside in mucosal tissues, allowing them to respond rapidly to stress signals without the need for traditional MHC presentation (Holtmeier & Kabelitz, 2005; J. Li et al., 2024). Understanding receptor-ligand specificity is essential for interpreting pathological immune responses, such as autoimmune reactions, just as it is for characterizing healthy immune function.

The human body generates a vast number of TCRs and BCRs, with the total number of unique receptors estimated to be around 10^9^ in the peripheral blood of a healthy individual (Greiff, Bhat, et al., 2015; Greiff, Miho, Menzel, & Reddy, 2015). On the antigen-binding region of each receptor are three Complementarity-determining regions (CDRs) numbered sequentially from 1 to 3, with CDR3 being the most variable and directly involved in antigen recognition **(Figure 1A)**. The remarkable diversity of BCRs and TCRs is achieved through somatic “VDJ” recombination, a genetic rearrangement that occurs in developing lymphocytes (Hozumi & Tonegawa, 1976). The core molecular events involve the RAG-1 and RAG-2 proteins recognizing and cleaving specific Recombination Signal Sequences (RSSs) flanking the gene segments, followed by the non-homologous end joining (NHEJ) pathway which repairs the DNA breaks, leading to the imprecise joining of the V, D, and J segments and the formation of the coding joint, thereby generating junctional diversity (Bassing, Swag, & Alt, 2002; Christie, Fijen, & Rothenberg, 2022). VDJ recombination occurs in the Beta (TRB) chain of αβ T cell lineages, and the Delta (TRD) chain of γδ lineages (Kotrova, Darzentas, Pott, Baldus, & Brüggemann, 2021). In B cells, recombination events are central to the generation of the immunoglobulin heavy (IGH) chain. Following antigen exposure, B cells additionally enhance their specificity through somatic hypermutation (SHM), a process by which high-frequency mutations are introduced to immunoglobulin genes to further refine their antigen binding capacity (Janeway, Travers, Walport, & Shlomchik, 2001b). In the case of αβ T cells, the MHC presentation and the vast number of class I and II alleles in human populations means antigen specificity of any T cell is a function of both the TCR and the MHC background (Klein, Kyewski, Allen, & Hogquist, 2014).

**Figure 1.**
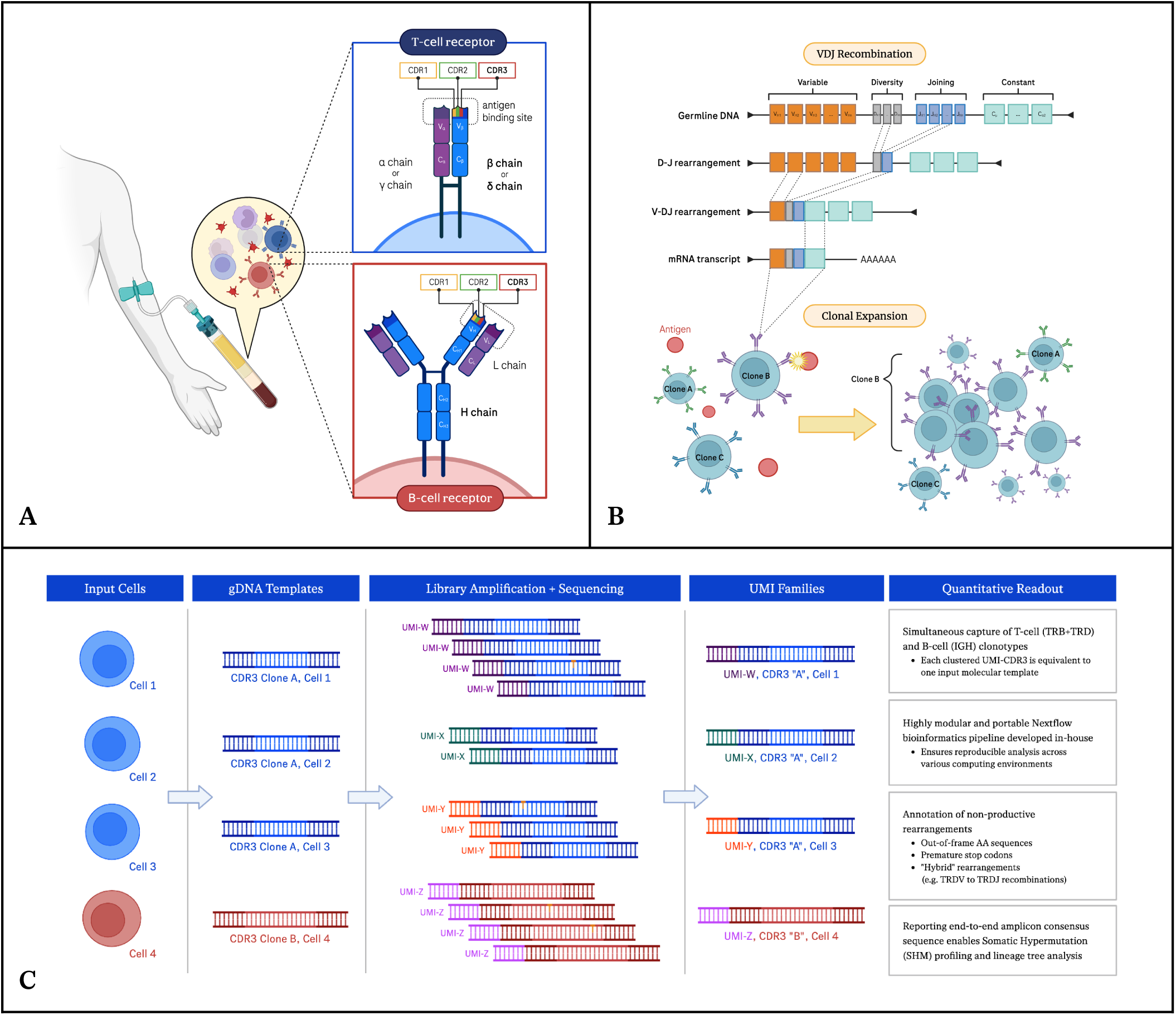
Schematic diagram of VDJ recombination and immunoPETE workflow. **A.** Structure of the T-cell receptor (TCR) and B-cell receptor (BCR) created with Biorender - TCRs and BCRs feature an antigen-binding site with three complementarity-determining regions (CDRs), with CDR3 being the most variable and critical for antigen recognition. **B.** VDJ recombination and clonal expansion - Variable, diversity, and joining genes of TRB, TRD, IGH chains are rearranged to create unique receptor sequences, generating immense potential diversity at the population level. Activated lymphocytes undergo clonal expansion, rapidly dividing to create a large population of cells with matching specificity. **C.** Genomic DNA is extracted from lymphocytes with recombined VDJ arrangements. UMI-based deduplication with PCR error correction enables quantifying of CDR3s in TCR and BCR subsets at a cell-level resolution. Next, an in-house data processing Nextflow pipeline called Daedalus generates sample readouts (CDR3-cell report, clone report) and an overall summary of repertoire-level metrics (e.g. read depth, cell recovery, CDR3 diversity).

When a lymphocyte’s unique receptor successfully binds to a specific antigen in the presence of co-stimulating signals, the cell (having previously passed developmental checkpoints such as negative selection) enters an activated state and undergoes clonal expansion **(Figure 1B).** During this phase, the lymphocyte rapidly divides, creating a large population of cells that share the same antigen specificity (sometimes with improved antigen binding through somatic hypermutation in B cells) (Goodnow, Sprent, de St Groth, & Vinuesa, 2005). Each collectively expanded group of cells descended from the same ancestral B or T cell is referred to as a clone. The collection of these unique clones and their abundance is known as the immune repertoire.

Immune repertoire profiling is the comprehensive study of adaptive immune receptor repertoire (AIRR). At the molecular level, immune repertoire profiling can be performed using either RNA or DNA as the starting template. RNA-based assays benefit from the higher copy number of mRNA per cell compared to the DNA template and may be more sensitive to rare clones. However, their focus on active transcription combined with the instability of mRNA limits its ability to capture the complete repertoire, and makes any quantitative measurement vulnerable to biases due to expression level variation of highly transcriptionally active cell populations (Trück et al., 2021). In contrast, DNA-based profiling, with its equivalence of one DNA template to one cell, provides a stable and comprehensive snapshot of receptor diversity by including both active and bystander lymphocyte populations (Mazzotti et al., 2022; Pai & Satpathy, 2021; Seo & Choi, 2025). This makes it ideal for applications requiring a complete mapping of the immune system’s architecture, such as tracking clonal evolution in cancer (Che et al., 2025).

This manuscript introduces immunoPETE, an assay that simultaneously targets recombined human TRB, TRD, and IGH chains from genomic DNA (gDNA) through a two-stage primer extension and enrichment process. Integration of Unique Molecular Identifier (UMI) onto each template molecule enables accurate quantitation of clonal composition at a single cell-level resolution. immunoPETE has been used to study samples from various input sources including whole blood, peripheral blood mononuclear cells (PBMCs), formalin-fixed paraffin-embedded (FFPE) tumor tissues, urine, in addition to some early investigative work on cerebrospinal fluid (CSF), generating valuable insights in the study of cancer, infectious diseases and autoimmune disorders (Akbari et al., 2025; Dannebaum et al., 2022; Joseph et al., 2022; Schneider-Hohendorf et al., 2022, 2025).

In this manuscript, we first describe the laboratory and computational workflows of the immunoPETE assay. Second, we report a comprehensive evaluation of the assay’s analytical performance. Third, we present a case study from a cohort of non-muscle-invasive bladder cancer (NMIBC) patients to demonstrate an example of immunoPETE’s application in oncology research. Finally, we summarize our findings, briefly describe selected examples of previously published studies using immunoPETE and discuss the potential applications and future directions of this technology. Note that immunoPETE is intended for Research Use Only (RUO), and not for use in diagnostic procedures.

## 2. immunoPETE: An assay for T-cell Receptor (TCR) and B-cell Receptor (BCR) Profiling

immunoPETE is a primer-extension-based targeted gene enrichment assay derived from Roche’s PETE technology ([patent US10731212B2]) (US Patent & Trademark Office Patent No. 10731212, 2019). The assay employs a two-stage multiplex PCR approach to target specific loci using genomic DNA (gDNA) as the input **(Figure S1)**. From gDNA, an initial primer extension is performed with a pool of V-gene oligos which contain an 11-base unique molecular identifier (UMI) with anchors and a universal amplification sequence. An exonuclease treatment followed by bead-based purification removes remaining oligos and unextended gDNA. Next, a master mix containing a pool of J-gene oligos, an i7-bridging primer and i7/i5-sequencing primers with dual unique indexes are added to the purified V gene-primed templates for J gene primer extension. The resulting libraries are purified on magnetic beads, then quantified and analyzed for fragment size. Libraries are combined in equal mass or volume to form a library pool which undergoes additional bead-based purification, quantification and fragment analysis prior to sequencing using Illumina sequencing platforms. Samples are typically sequenced on a dedicated flow cell using 2×150-bp paired-end reads.

Raw demultiplexed FASTQs are assessed for conformance in UMI patterns, followed by optional downsampling of valid reads to reduce computational burden. Read mate pairs are merged using FLASH (Magoč & Salzberg, 2011), trimmed of primer templates, and aligned to IG and TR references from ENSEMBL annotated in accordance with the international ImMunoGeneTics information system (IMGT) nomenclature (Lefranc, 2014). High quality gene-aligned reads are clustered into families based on the combination of CDR3 and UMI identities, which we will refer to as “UMI families” **(Figure 1C).** UMI-based clustering of reads enables error correction and generation of consensus sequences in the case of discordant base calls among reads within a family. UMI families stemming from potential cross-contamination among samples are removed using a proprietary algorithm, and functional arrangements are identified based on the presence of non-chimeric, non-pseudogene V/J-gene annotations, a minimum CDR3 length of 3 amino acids (AAs) and the absence of ambiguous AAs, stop codons, and frameshift mutations in the CDR3 sequence.

All CDR3s are reported for each processed library, where the number of functional (productive) rearrangements corresponds to the count of viable cells that were recovered. V/J-gene annotations, CDR3 sequence, functional classification, and full-amplicon consensus are reported for each captured rearrangement. The standard pipeline also reports repertoire-level QC summaries consisting of run metrics and diversity measures to provide a high-level overview of the experiment. **Table S1** describes the content of reports generated by the immunoPETE assay. Individual samples pass QC if all of the following three conditions are met: 1) The pipeline successfully completes all data processing steps and generates the standard reports, 2) 50000 or more reads are recovered, and 3) 10 cells or more are recovered. Conditions 2 and 3 were relaxed in this study for serial dilution samples where low input DNA masses were used by design.

## 3. Assay Performance Characterization

The immunoPETE assay was evaluated through a series of experiments designed to assess its accuracy, reproducibility, sensitivity, and versatility of usage. Samples consisting of contrived mixtures of known lymphocyte populations were used to evaluate the accuracy and precision of clonal profiles. Quantitative metrics such as the Limit of Blank (LoB), Limit of Detection (LoD), and Limit of Quantitation (LoQ) were measured using high-complexity PBMCs from multiple donors. We also assessed the impact of input DNA quantity, sequencing platform, sequencing depth, and read subsampling to optimize cell recovery. The ability to detect rare clones was assessed using a HuT-78 T-lymphocyte cell line spiked into PBMC background at serially decreasing fractions. All samples were processed in replicates. “Replicates” in this experiment were created by extracting aliquots from the prepared DNA (extracted or mixed; before any amplification) and running immunoPETE on them independently. Therefore, each replicate contained a unique set of template DNA molecules. A schematic summary of the experiment is presented in **Figure 2**.

**Figure 2.**
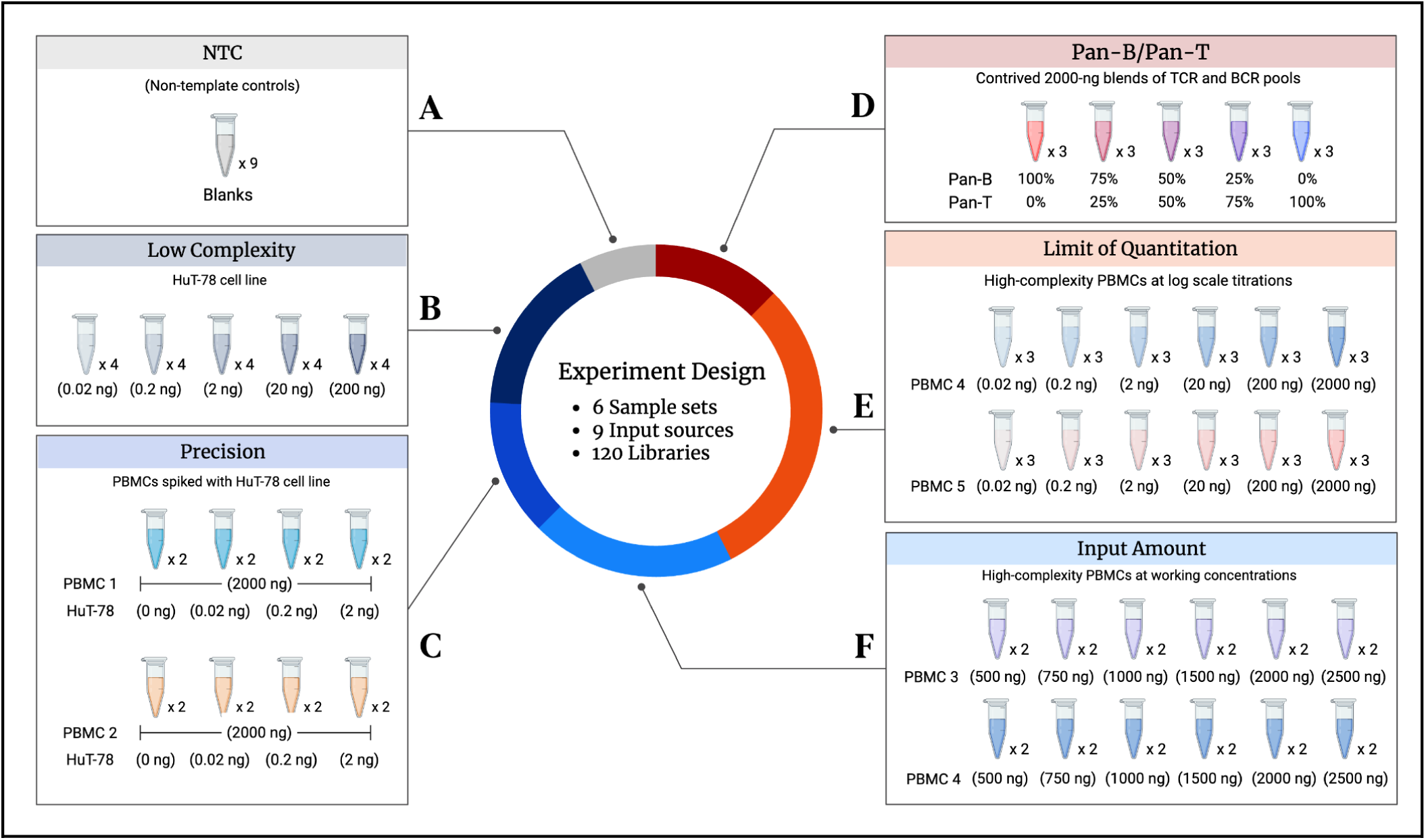
Schematic overview of the immunoPETE analytical performance experimental design. The study evaluated assay accuracy, reproducibility, and sensitivity using 120 independent libraries allocated across 6 distinct sample sets. **A.** NTC: No-template controls were used to determine Limit of Blank (LOB) and measure baseline contamination levels. **B.** Low-Complexity: Five log-fold titrations of the HuT-78 monoclonal cell line were designed to estimate the range of linearity and UMI collision limits in low-complexity libraries. **C.** Precision: Two PBMC sources (PBMC1, PBMC2) each distributed into eight 2000-ng libraries were used to quantify precision (reproducibility) of clonal compositions and other metrics in high-complexity libraries. They were spiked with log-increments of the HuT-78 cell line in duplicates to assess the recovery of rare clonotypes*. **D.** Pan-B/Pan-T: Contrived mixtures of DNA extracted from viable T cell and B cell stocks at set ratios enable retrieval of cell type ratios (i.e. %TRB, %TRD, and %IGH) and V-gene and J-gene usage frequencies*. **E.** Limit of Quantitation (LoQ): Log-fold increments of PBMCs from two additional sources (PBMC4, PBMC5) were designed to determine the limits of quantitation, assess sample linearity on a log scale from 0.02 ng to 2000 ng*. **F.** Input Amount: Two PBMC sources (PBMC3, PBMC4) were sequenced at six input levels within the expected working range to estimate linearity of yield and determine optimal input mass. *All libraries in the Pan-B/Pan-T mixtures and LOQ set include a constant 2-ng background spike-in of the HuT-78 cell line. Hut-78 sequences were bioinformatically removed from these and the precision set repertoires before PBMC-specific analyses for reproducibility, linearity and cell type ratio retrieval.

A total of 120 libraries were prepared from a diverse set of samples, including negative controls, samples from five unique PBMC stocks (Stemcell Technologies CAT #70025), the HuT-78 cell line (RMSCC CAT #0011147), and purified T-cell (Pan-T) and B-cell (Pan-B) mixtures (Stemcell Technologies Cat #70023 - 70024). Individual libraries were strategically distributed across five processing batches, each with its dedicated flow-cell in order to detect and quantify batch effects. The same libraries were subsequently pooled into a single flow-cell and sequenced in one batch on the NovaSeq platform. **Table S2** describes the detailed experimental design of the analytical performance characterization study. Certain high complexity samples were spiked with HuT-78 cells **(Table S2)**. To evaluate the power to detect rare clones, we analyzed the recovery of HuT-78 cells in the “precision set” samples, specifically at the 10^-3^, 10^-4^, 10^-5^ and 0 fractions in a 2000 ng PBMC background. Cell line sequences were bioinformatically removed from these high complexity repertoires prior to other (PBMC-focused) analyses.

### 3.1. Assay fidelity and reproducibility

Accuracy (fidelity) is the capacity of an assay to accurately retrieve the ground truth whereas reproducibility (precision) quantifies the similarity of measurements across multiple replicates of the same sample. Our analysis centers on three key features from the immunoPETE output: CDR3 sequence and clone identity, B/T cell type fraction, and total cell count. However, accuracy and reproducibility are in principle extractable for any of the features generated by immunoPETE.

#### 3.1.1. CDR3 sequence accuracy and clone identity

We evaluated the assay’s sequence retrieval accuracy and reproducibility using the HuT-78 cell line, a well-characterized monoclonal T-cell line. Cell-line libraries were sequenced in quadruplicates across DNA inputs ranging in log-fold increments from 0.02 ng to 200 ng. For each library, we determined the percentage of cells that exhibit exact V-gene and J-gene annotations, and a CDR3 (nt) sequence above 95% homology to the reference in **Table 1**.

**Table 1.**
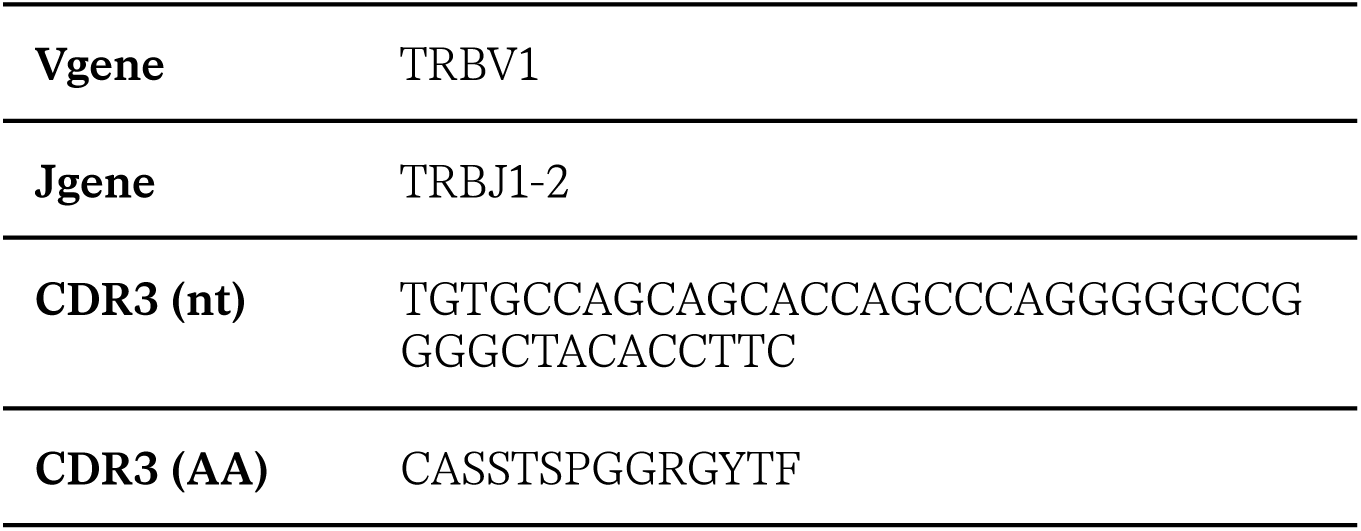
HuT-78 Cell line clonotype ID.

Replicates with a DNA input of 2 ng or more consistently showed purity > 98% in this experiment **(Figure 3A)**. The observed sequence impurity in low-input samples likely stems from cross-sample contamination, rather than mutations, experimental artifacts, bioinformatic errors, or other sources. This is supported by two key findings. First, most of the unexpected sequences significantly deviated from the canonical HuT-78 sequence, suggesting external origin. Second, both the mean and variance of sequence impurity decreased with higher input amounts. If other factors were at play, the average sequence accuracy would be similarly affected across different input levels. This pattern indicates a low level of read artifacts that is prominent at very low input levels but becomes negligible at input amounts typically used in real experiments (99.9% purity at 200ng).

**Figure 3.**
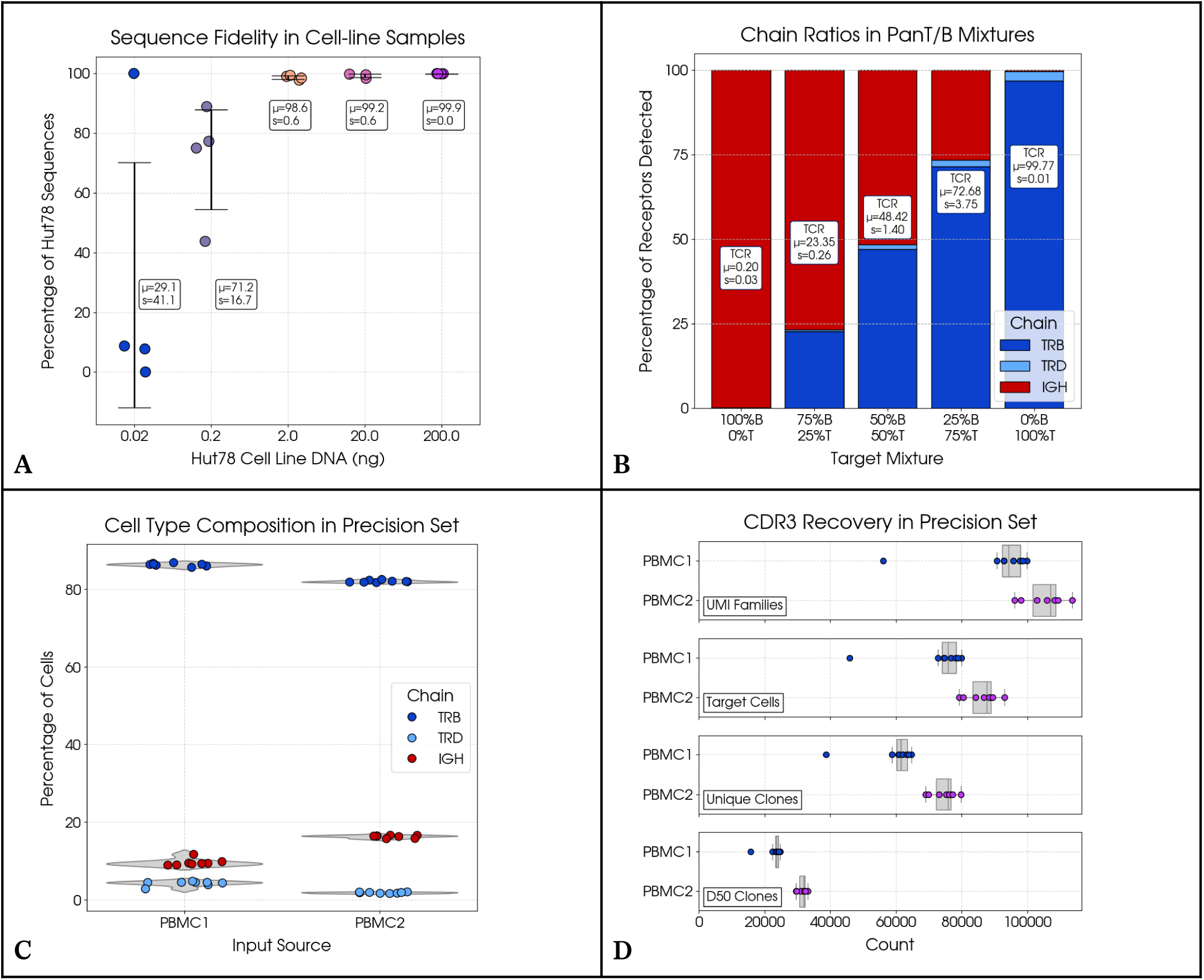
Evaluation of accuracy and precision of clone identity, cell count and cell-type composition. **A.** Observed fraction of the canonical HuT-78 clone sequence across varying DNA input amounts from 0.2 ng to 200 ng - At inputs above 2 ng, clonal purity was consistently >98% with minimal variation across replicates. **B.** Quantification of IGH and TRB fractions in contrived Pan-T and Pan-B mixes - The assay accurately detected expected T-cell and B-cell ratios across mixtures, with samples reflecting anticipated IGH percentages with low variability. **C.** Cell type composition in precision set - Violin plots show the distribution of chain fractions from the 8 replicates of PBMC1 and PBMC2. Reproducible fractions within each input source demonstrates the high precision of the cell type composition measurement. **D.** Quantification of CDR3 recovery - PBMC2 observes higher recovery of unique VDJ-rearranged templates (UMI families) compared to PBMC1, resulting in higher target cell count (TCR+BCR) and unique clonotypes.

#### 3.1.2. Recovery of T-cell and B-cell fractions in contrived mixtures

immunoPETE enables accurate simultaneous capture of T-cell and B-cell subsets, a feature that is often lacking in traditional RNA-based repertoire sequencing assays. In order to evaluate the recovery of original T-cell and B-cell proportions, we prepared Pan-T cells (pure T-cell populations isolated from human peripheral blood), Pan-B cells, and synthetic Pan-T/Pan-B mixtures at increments of 25% to a total DNA input mass of 2000 ng per library with three replicates per mixture **(Figure 3B).** Annotated cell counts of the Pan-T/Pan-B mixtures recovered from the assay consistently matched targeted ratios with the Mean Absolute Error ≤ 3% of the targeted ratio (of the major component), and Coefficient of Variation (CV) ≤ 0.15 for BCRs and TCRs in all mixtures **(Table S3).**

#### 3.1.3. Reproducibility of key repertoire metrics in high-complexity samples

The precision set of samples consists of two sources (PBMC1, PBMC2) of high-complexity PBMCs supplied by Stemcell Technologies. We sequenced 2000 ng of input DNA with 8 technical replicates per source to assess the assay reproducibility through the consistency of key repertoire metrics, namely cell recovery, chain fractions, and repertoire diversity scores **(Table S4).**

Both samples in the precision set exhibited predominant T-cell populations (TRB and TRD chains), with B-cells (IGH) comprising a smaller fraction, though PBMC2 had a slightly higher relative representation of IGH. Again, we consistently observed steady cell type ratios, with recorded CV ≤ 0.15 for all three chains in both PBMCs **(Figure 3C).** We also observed that recovery of unique VDJ rearrangements were similar across replicates of the same donor, with PBMC2 showing a higher number of UMI families from the 2000 ng of starting input, which was carried forward into cell counts and unique clonotypes **(Figure 3D).** The observed variation in cell recovery yield (total cell count per input ng DNA) among individual donors, as seen here and in later analysis, is due to several factors including the fraction of lymphocytes (immunoPETE target cells) in their PBMC. One out of the eight PBMC1 replicates yielded a smaller number of cells than others, resulting in a higher cell count CV (PBMC1: CV = 0.15 vs. PBMC2: CV = 0.05). Despite the lower absolute cell recovery, the clonal composition was preserved in this replicate - which is important as we examine repertoire architecture (see section 3.1.4).

The clonal space diversity can be characterized by the aggregate frequency of top clones. For example, the D50 score represents the number of unique clonotypes occupying 50% of all recovered cells in the library. 38.7% ± 0.9% (mean ± SD) of clones in PBMC1 are in the top 50% of clonal space, indicating a higher proportion of dominant clones compared to 42.2% ± 0.3% of clones in PBMC2. We further investigated the reproducibility of top clone contribution across replicates by extracting the most dominant (n=10) clones in each donor, and determined that the cumulative abundance of the ten clones account for 3.92% ± 0.19% of clonal space in PBMC1 and 1.31% ± 0.04% in PBMC2 **(Table S5).**

#### 3.1.4. Reproducibility of dominant clone contributions

In this section, we assess the intrinsic characteristics of the repertoire to ensure integrity in the recovery of TCR and BCR clones. Following the observation that the key summary statistics, e.g. cell counts and cell type ratios, were consistent among replicates, we proceeded to examine the identity of top clones across replicates and their abundance. We again utilized the precision set consisting of two PBMC donors sequenced at 2,000 ng in eight replicates. The relative abundance of the top 10 clones was highly reproducible across all replicates of the same input source, despite slight variations in total cell count **(Figure 4A).** Crucially, this high reproducibility persisted even for clones comprising a minor fraction of the repertoire (e.g. <1% of all cells), with CV ranging from 0.03 to 0.27 in PBMC1, and from 0.07 to 0.20 in PBMC2 **(Table S5).**

**Figure 4.**
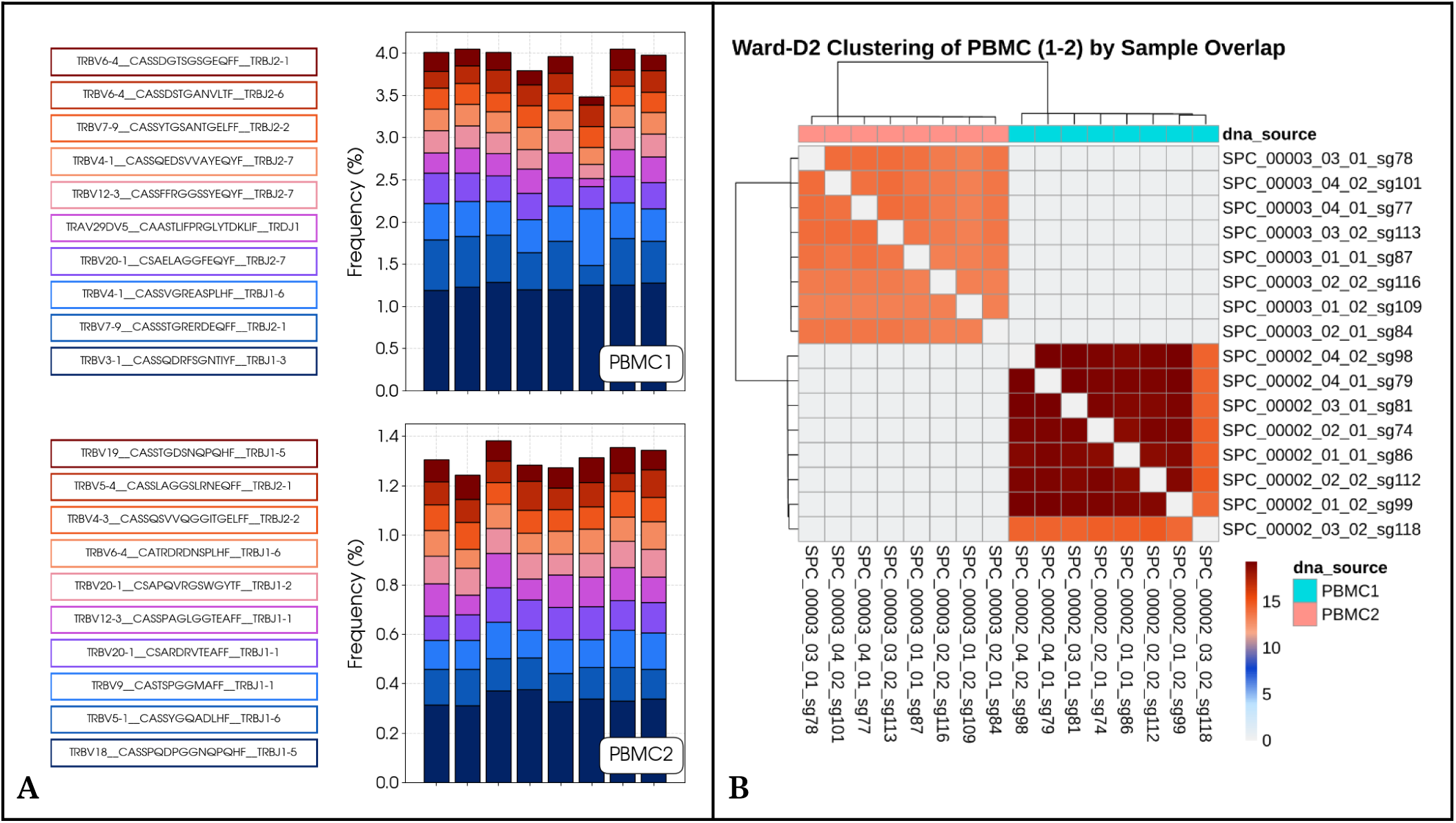
Profiling of top clones and repertoire similarity in the precision set (PBMC1, PBMC2) **A.** Stacked bar plots show the frequency for each of the top 10 most prevalent clones in PBMC1 (top) and PBMC2 (bottom), sequenced with 8 replicates at 2000 ng of gDNA per replicate. The total cell recovery in each library is also displayed. All clones were recovered at similar abundance within a PBMC donor, despite variations in cell count **B.** Pairwise similarity for PBMC1 and PBMC2 libraries was represented by the Sørensen similarity index (Equation 2), characterized by the percentage (S × 100%) of aggregated shared cell counts between two samples belonging to the same clonotype. Libraries that share similar overlap patterns were clustered using the Ward-D2 method. See Table S2 for information on specific replicate description and attributes.

#### 3.1.5. Relationship between pairwise similarity and repertoire diversity

Pairwise similarities between all replicates from two individual PBMC donors (PBMC1, PBMC2) were calculated using the formula for Sørensen similarity index (S) shown in **Equation 1** (Sørensen, 1948). Samples were clustered using the Ward-D2 method based on the similarity matrix using the *pheatmap* package in R **(Figure 4B)**. Replicates of each donor clustered together with a normalized S index in the 0.13-0.20 range. Replicates from the two different donors showed minimal overlap (S < 0.01) leading to perfect segregation of two clusters corresponding to each specific donor. Average overlap was higher among PBMC1 replicates compared with PBMC2 replicates. We hypothesized that the extent of overlap between replicates is influenced by the overall diversity of the repertoire. Highly clonal repertoires are expected to exhibit greater overlap between replicates compared to highly diverse repertoires. This is because there is an increased probability of detecting DNA from cells belonging to high-frequency clones in independent replicates. Consistent with this assumption, PBMC1 had a larger Gini index than PBMC2 (PBMC1: 0.181 ± 0.012, PBMC2: 0.132 ± 0.005) **(Table S4).** Gini Index (G) is a measure of compositional inequality where higher values closer to 1 indicate a repertoire that is dominated by fewer clones.

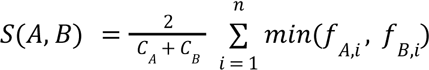

**Equation 1**. Sørensen similarity index

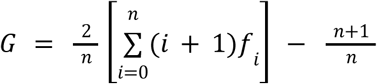

**Equation 2**. Normalized Gini index

*f_A,i_*, *f_B,i_*: Abundance of clone i in samples A and B

*C_A_*, *C_B_*: Total number of cells in samples A and B

*n*: Total number of cells in samples A and B

#### 3.1.6. Reproducibility of clonal composition at different DNA input masses

Having demonstrated the consistency in profiling top clones within the fixed-input precision set, we next sought to emphasize assay robustness across a dynamic working range of input mass. For this analysis, the input amount set (PBMC3, PBMC4) was processed, consisting of gDNA amounts ranging from 500 ng to 2,500 ng run in duplicate at each level. Three PBMC3 libraries were excluded due to failed quality control. From the passing libraries, we confirmed that the relative abundance of the top clones remained highly reproducible within each input source and consistent across the entire 500 ng to 2,500 ng range of gDNA input **(Figure S2).** The reproducibility of the abundance of top clones was measured by CVs ranging from 0.10 to 0.20 in PBMC3, and 0.04 to 0.14 in PBMC4 **(Table S5).** Similarly, individual replicates were clustered using the Ward-D2 method. While replicates within the sample showed similar overlap profiles irrespective of input mass, the absolute degree of similarity was modest (S ≤ 0.20) - consistent with results from the precision set.

Overall, our analysis showed that immune repertoire readouts remain consistent across replicates (aliquots) from the same sample, allowing perfect clustering of samples based on the donor, regardless of input mass. This observation signifies the assay’s reliability to capture a representative, individually distinct sample of the immune repertoire despite some variation in sample quality or low gDNA input amounts in real-world applications.

#### 3.1.7. Concordance of V-gene and J-gene usage profiles in Pan-T / Pan-B mixtures

Building upon the initial characterization of cell type ratios in contrived Pan-T/Pan-B mixtures, we proceeded to analyze the resulting V-gene and J-gene usage frequencies in each mixture. Within replicates of the same mixture, the relative proportions of V-genes and J-genes are strongly correlated, suggesting a reproducible representation and indicating that the small fraction of adaptive immune cells sampled in the blood is representative of the repertoire’s overall V/J gene usage profile (**Figure 5A).** Comparing Pan-B to a 50% Pan-T / 50% Pan-B mixture shows a shift, with IGHV relative gene frequencies remaining largely unchanged despite the expectedly increased frequency of TCR populations (TRBV and TRDV genes). A similar effect was observed for TRBV genes in Pan-T compared to the same Pan-T / Pan-B mixture. Pan-B and Pan-T samples were only guaranteed by the manufacturer to be ≥85% pure based on detected CD19+ and CD3+ markers through flow cytometry, respectively (STEMCELL Technologies, 2021a, STEMCELL Technologies, 2021b), thus explaining the non-zero detection of TCRs in Pan-B mixtures and BCRs in Pan-T mixtures. The relative hierarchy of TRBV genes between the Pan-B replicates and the relative hierarchy of IGHV genes between the Pan-T replicates were still well-preserved suggesting that even at a low residual input level, the assay reflected repertoire composition. Note that although gene frequencies are usually calculated and normalized within each class (IGH, TRB, TRD) separately, we chose to combine and normalize them within a single pool in this analysis to capture cell type fractions and gene frequencies in one plot for each comparison.

**Figure 5.**
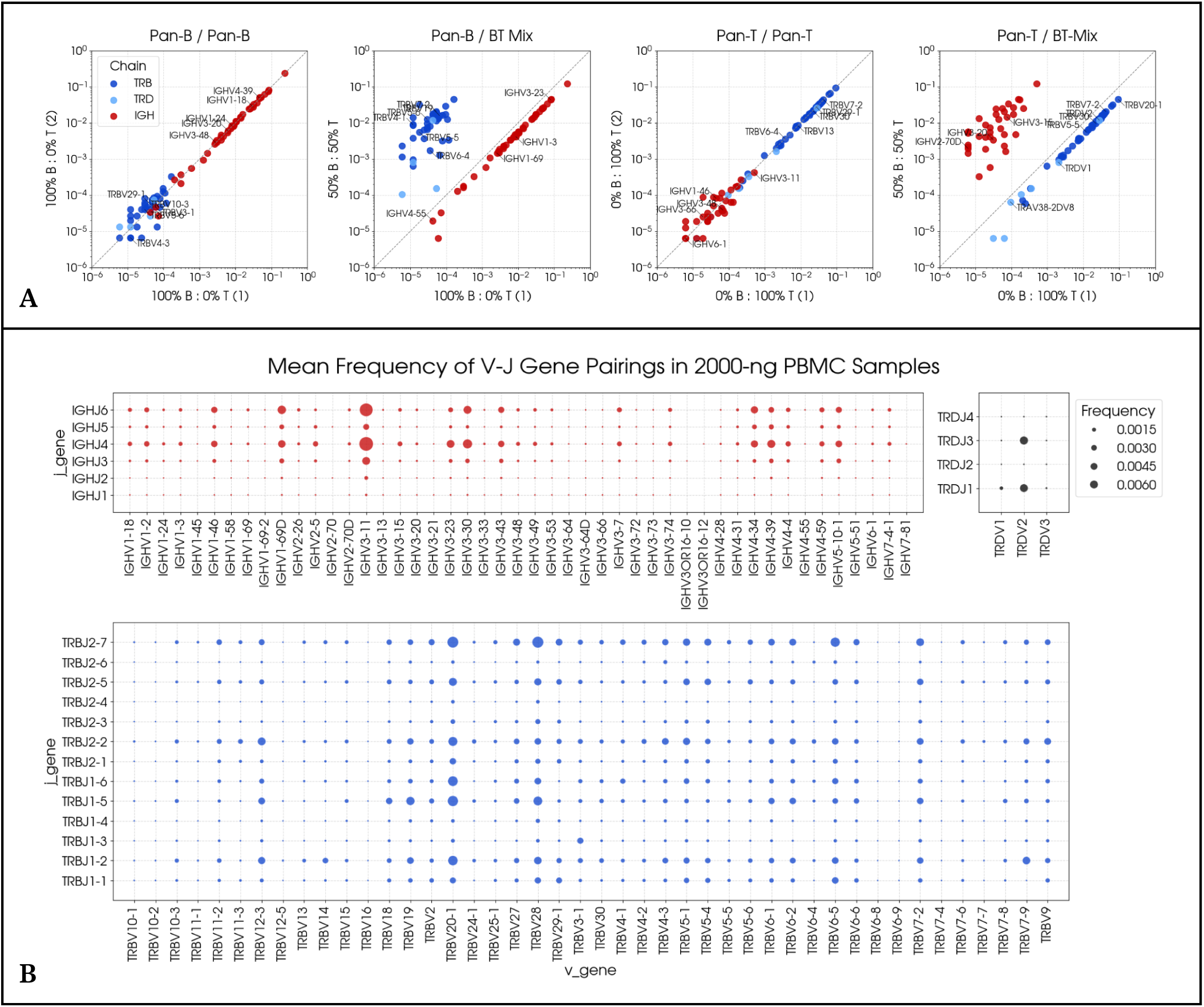
Characterizing repertoires by CDR3 composition and V/J gene usage. **A.** Within technical replicates of the same pure mixture (Pan-B / Pan-B and Pan-T / Pan-T), the relative proportions of V-genes and J-genes are highly concordant. This demonstrates technical reproducibility and low sampling variability. Transitioning from the pure BCR replicates to mixed BT mixture results in a diminishment of IGHV genes and the concurrent emergence of TRBV/TRDV genes. A similar effect was observed when shifting from the pure pan-T replicates to the BT mixture. **B.** Mean frequency of V/J gene pairings in five PBMC samples. Aggregated cell counts from two replicates per PBMC sample at the 2000 ng input level were normalized to relative frequencies. Each point represents a V/J gene pairing, with the point size indicating the abundance of associated cells.

To further evaluate how the gene usage profiles shift across a range of TCR and BCR compositions, aggregated V-gene and J-gene frequencies were determined for all three replicates within each B/T ratio class of mixtures. The pairwise scatterplot matrix in **Figure S3** compares V-gene and J-gene profiles between all mixtures (the two replicates from each biosample were aggregated for this analysis). Consistent with previous results, our analysis demonstrates a higher degree of concordance in gene frequencies between mixtures of similar composition. The relationship shifts clearly with the targeted input ratios, with IGH genes (red) showing dominance in B-cell rich mixtures, and TCR genes (blue) showing increasing prominence as T-cell fraction increases, confirming the distinct recovery of both repertoire types.

#### 3.1.8. Repertoire fingerprinting with concurrent V/J gene usage frequencies

After examining individual V and J gene frequencies, we explored the concurrent usage (pairing) of specific V and J genes. First, to illustrate high-level patterns among all PBMC samples in this study, cell counts from the five PBMC samples (PBMC1-5) were aggregated based on the V/J gene usage. To ensure equal representation of all inputs, aggregation was performed on two replicates per PBMC donor at the 2000-ng input level, and the resulting cell counts were normalized into relative abundance. **Figure 5B** presents the mean frequency of each cassette in a bubble plot. This visualization highlights prominent gene pairings and their relative distributions across the dataset. Recent evidence challenges the traditional view that the selection of V and J genes during VDJ recombination occurs independently, particularly regarding the TRB chain. (Levi & Louzoun, 2022) Given immunoPETE’s ability to quantify yield at a cell level, we combined the exact cell count for all existing V and J cassette combinations using two 2000-ng replicates from each of the five donors (PBMC1-5). To understand whether V-J pairings in our dataset exhibit systematic bias, we performed a Chi-squared (IZ^2^) test of independence on TRB, TRD, and IGH counts from each donor. Every test yields a highly significant result (p-value≤1E-32) with Cramér’s V values in 0.077-0.201 range, confirming that pairing of V and J segments is partly non-random in every analyzed immune receptor repertoire **(Table S6).** Cramér’s V (ϕ_*c*_) statistic measures the effect size for association of nominal variables calculated from ^2^ tests of independence with a value of 0 indicating complete independence (no association) and 1 indicating complete dependence (when the value of one variable can be perfectly predicted by the other variable). The TRD chain consistently shows the highest value across all donors, suggesting the strongest degree of non-random V-J pairing, followed by TRB and IGH.

To further explore gene usage trends, all replicate libraries from PBMCs with a minimum of 3 replicates at 2000 ng input were included. Gene usage was calculated at the replicate level and z-transformation was applied to standardize the data. t-distributed Stochastic Neighbor Embedding (t-SNE) (Maaten & Hinton, 2008) was then applied using the scikit-learn library in Python, with a perplexity value of 15, enabling the identification of distinct sample-specific clusters **(Figure S4).** Although not easily generalizable to a larger number of samples with varying degrees of diversity and similarity, this exploratory analysis exemplifies the potential of gene frequencies in sample identification and classification.

### 3.2. Quantitative limits of immunoPETE

General assay performance metrics such as limit of blank (LoB), limit of detection (LoD) and limit of quantitation (LoQ) must be interpreted according to the nature of the assay under evaluation. In principle, they can be calculated for any of the multiple quantitative measures extracted from immune repertoire data. Here, we focus on two instances that are conceptually important and relevant to specific downstream applications: quantitative accuracy of total cell count, and the ability to detect specific clones at low frequency.

#### 3.2.1. LoB, LoD and LoQ of total cell count

The quantitative range of an assay is typically established by the span over which both acceptable linearity between the input and output and sufficient replicate reproducibility (precision) are achieved. We evaluated the linearity of total detected cells vs the input DNA mass in both low complexity (cell line) and high complexity (PBMC) samples. Linearity in high-complexity samples was assessed in two ways: across a wide range, including lower concentrations, using a logarithmic scale, and within a practical range (500–2500 ng) with 500 ng intervals, representing realistic expected conditions **(Figure 6).** In general, higher DNA input masses can lead to reduced recovery efficiency due to factors such as primer depletion and polymerase competition amongst other PCR inhibitory effects. However, saturation effects were not observed up to 200ng in the low-complexity set, nor up to 2500 ng input in the high-complexity set.

**Figure 6.**
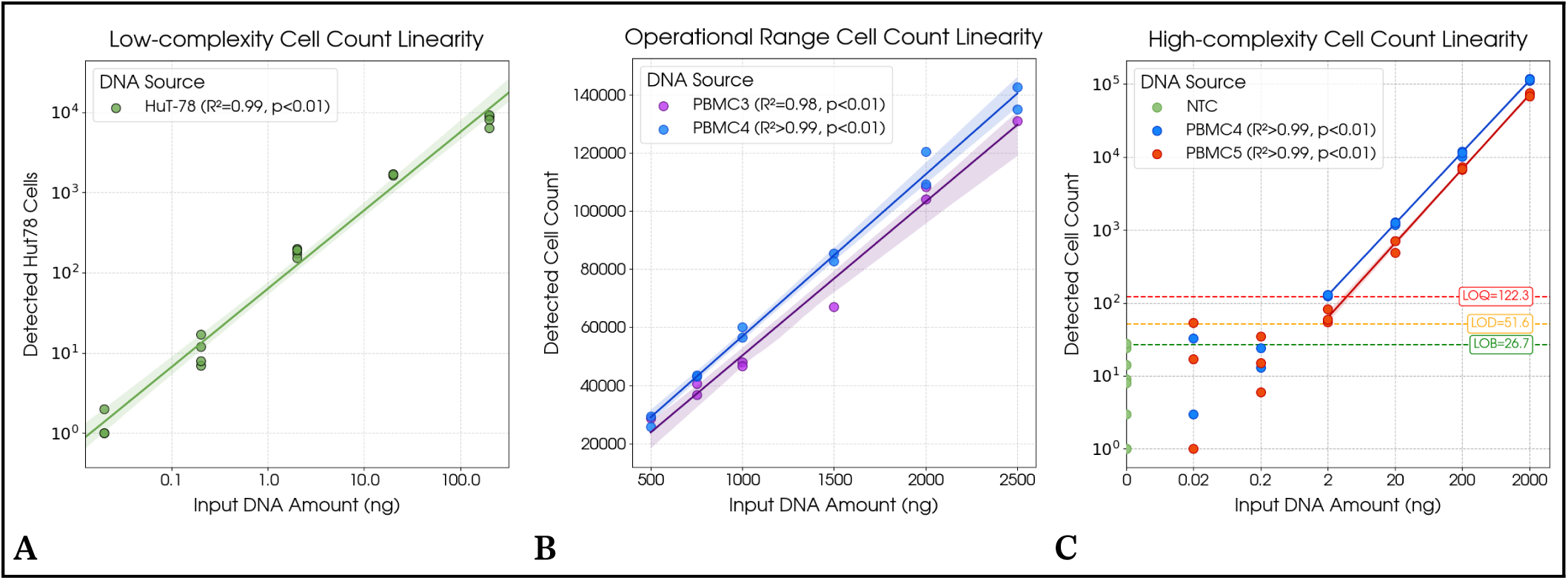
Input mass to cell count relationships in low-complexity cell line samples and high-complexity PBMCs. **A.** Strong linear correlation (R^2^ = 0.986) between cell recovery and DNA input mass in the low-complexity HuT-78 cell line, with no observed saturation effects (0.2–200 ng). **B.** Cell recovery exhibits strong linearity (R^2^ > 0.98) with DNA input mass within the standard working range (500 ng to 2500 ng) for the immunoPETE assay. High-complexity PBMC3 and PBMC4 samples demonstrate highly proportional cell recovery, confirming assay performance in the standard working window. The yield is slightly higher for PBMC4 across the range. **C.** The titration set spanning five orders of magnitude of input mass on the log scale (0.02 ng to 2000 ng) in two high-complexity samples (PBMC4 and PBMC5) establishes detection limits for the assay’s broad dynamic range and low-input sensitivity.

The standard operational input range was evaluated using DNA input masses from 500 ng to 2500 ng, which are typically expected by the immunoPETE assay for high-complexity samples such as PBMCs **(Figure 6B)**. Using PBMC3 and PBMC4, cell recovery was found to be highly proportional to input mass, demonstrating remarkably high coefficients of determination of R^2^ = 0.984 and R^2^ = 0.993, respectively. Our analysis confirms that the assay accurately and reliably reports the relative cell counts across this critical working window, confirming no primer depletion or saturation effects which can compromise cell recovery.

Complementing our findings from the linearity set at the operation range of inputs, we also observe a strong linear correlation between 2.0 ng and 2000 ng from the titration set (PBMC4, PBMC5). To quantify this relationship, the DNA input mass and detected cell counts were log-transformed, and we calculated the R^2^ value of the linear fit, yielding values of 0.999 and 0.998 for PBMC4 and PBMC5, respectively **(Figure 6C).** We excluded samples with input 0.02-0.2 ng from the linearity test due to low cell count reproducibility among replicates.

The relationship between input mass and cell recovery serves as a basis for establishing metrics related to detection limits, thereby improving the identification and quantification of low-abundance target cells with precision **(Figure 6C)**. The Limit of Blank (LoB) is defined as the number of recovered cells equal to the 95% upper limit in the no-template controls (NTCs). The limit of 26.7 cells as the LoB also serves as the upper limit for contamination, as detected cells within blanks are assumed to have originated from other libraries. The Limit of Detection (LoD) is defined equal to the minimum number of cells in a library that can be distinguished from NTCs with 95% confidence. The Limit of Quantitation (LoQ) is the lowest concentration within the range of input-output linearity with adequate precision. These measures can vary from sample to sample. In accordance with the Clinical and Laboratory Standards Institute (CLSI) guidelines, when testing two samples, we report the highest observed value for both the Limit of Detection (LoD = 51.6 cells) and the Limit of Quantitation (LoQ = 122.3 cells) from the two PBMCs.

#### 3.2.2. Detection of rare clones

To ensure the immunoPETE assay possessed the necessary sensitivity for detecting clinically relevant, low-frequency clonotypes, we designed a spike-in experiment where HuT-78 cell lines were added to 2000 ng of high-complexity samples PBMC1 and PBMC2 (the precision set). Our experiment utilized a titration series of HuT-78 cells spanning a log-scale concentration range to empirically determine the assay’s ability to extract rare clones. Analysis of the titration series confirmed that the immunoPETE assay detects the presence of the spiked HuT-78 clonotype with 100% sensitivity at 10^-3^ and 10^-4^ fractions using a single replicate with 2000 ng input in both PBMC1 and PBMC2 backgrounds. Sensitivity was 75% at the lowest tested fraction of 10^-5^ **(Figure 7).** Sensitivity can be improved by increasing the total amount of processed DNA (for example by increasing the number of replicates). A larger experiment will be needed in the future for more comprehensive evaluation of immunoPETE’s rare clone detection capabilities.

**Figure 7.**
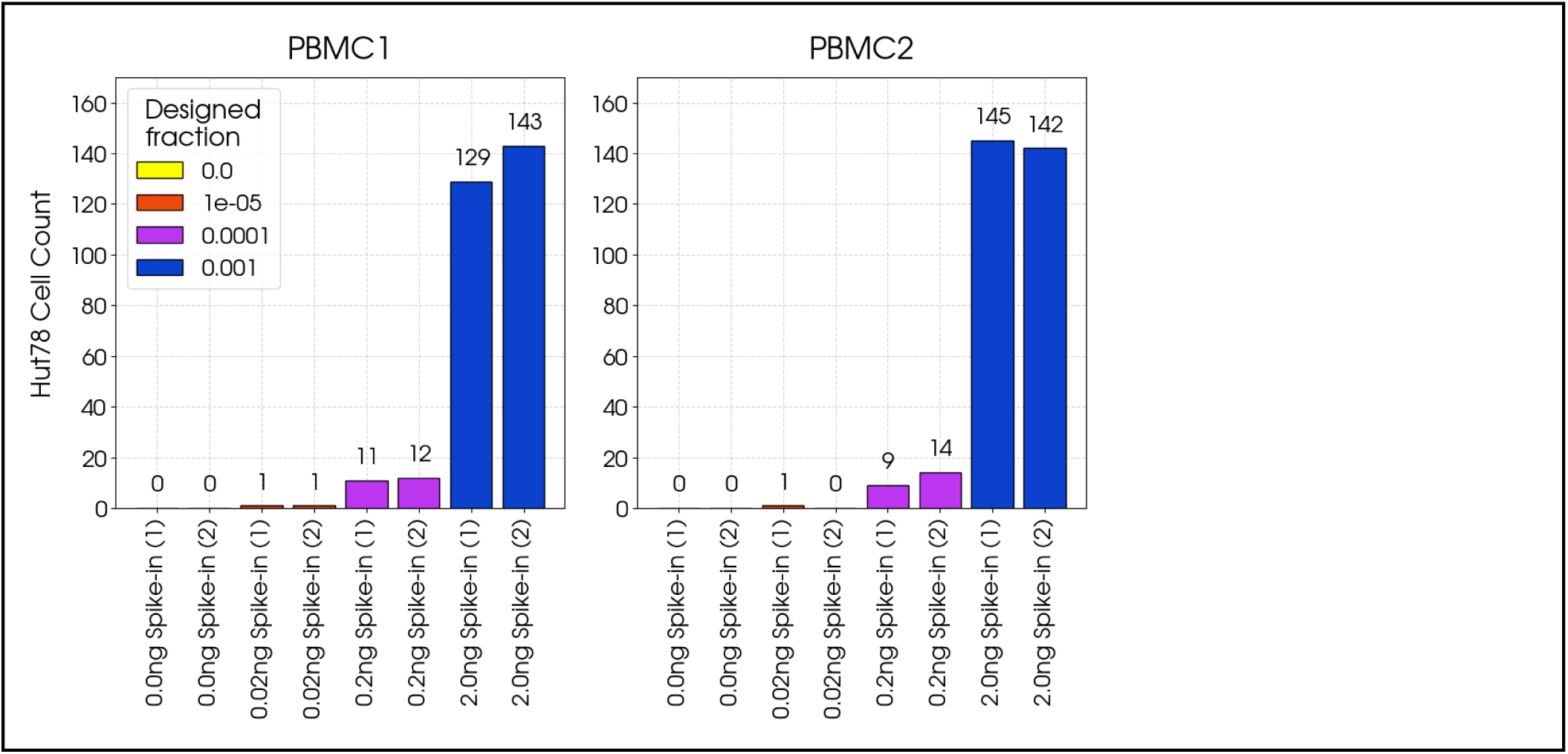
Recovery of HuT-78 spike-ins at ultra-low concentrations. Detection of HuT-78 cells spiked at varying concentrations into high-complexity PBMC samples. HuT78 cells were spiked into two 2000 ng PBMC background samples (PBMC1 and PBMC2) at designed fractions corresponding to 0.0 ng, 0.02 ng, 0.2 ng, and 2.0 ng of gDNA input, with two technical replicates for each condition. The assay detected the presence of the spiked clonotype down to the 0.02 ng input level (Designed fraction 10^-5^) across both PBMC backgrounds, demonstrating the sensitivity required for identifying rare clones.

## 4. Case Study: Immune Profiling in Non-Muscle-Invasive Bladder Cancer (NMIBC)^1^

To evaluate the utility of the immunoPETE assay, we conducted a pilot study on a cohort of NIMBC (Non-Muscle-Invasive Bladder Cancer) samples available through the West Midland’s Bladder Cancer Prognosis Programme where mature clinical outcome data was available (Zeegers et al., 2010). Paired fresh-frozen tumor and peripheral blood derived gDNA (collected pre-treatment) was available for 24 patients with non-muscle invasive bladder cancer (NMIBC)^2^ with 12 subjects also having urine-pellet gDNA available (ethics approval 06/MRE04/65). The majority of these patients (23/24) had high-risk NMIBC and subsequently received intravesical Bacillus Calmette-Guérin (BCG) therapy **(Table S7)**. All samples were processed in duplicate using the standard immunoPETE protocol to profile both B-cell and T-cell repertoires, with a specific focus on the often-overlooked Gamma/Delta (γδ) T cells.

This study aimed to address three key proof-of-concept questions: i) Can B and T cells be successfully recovered from urine samples? ii) Does the urine immune profile reflect that of the tumor or peripheral blood? iii) Can immune repertoire metrics derived from the immunoPETE assay serve as prognostic biomarkers for NMIBC patient outcomes?

### 4.1. Immune cell profiling from urine

Our findings confirm that in addition to PBMC, the immunoPETE assay can recover and profile immune repertoire profiles from urine pellets, a sample type often considered paucicellular in healthy individuals and mainly composed of shed epithelial cells from the bladder lining. (El Bali, Diman, Bernard, Roosens, & De Keersmaecker, 2014). For a single reaction, a median of 544ng (IQR 405ng to 692ng) of blood-derived gDNA, 464ng (IQR 374ng to 580ng) of tumor derived gDNA, or 379ng (IQR 147ng to 611ng) of urine cell pellet derived gDNA were used to prepare replicate libraries. While the number of B and T cells in urine was expectedly lower than in blood or tissue **(Figure 8A)**, the assay successfully generated repertoire libraries from all three specimen types analyzed. We observed a consistent trend in the relative fraction of cell types across all sample types (TRB > IGH > TRD) as shown in **Figure 8B**. Interestingly, the fraction of TRB and TRD chains was notably higher in tumor tissue and urine than in blood, suggesting a more significant infiltration of T cells into bladder tumors compared to B cells **(Figure 8C)**. Furthermore, the B cells infiltrating the tumor exhibited a significantly reduced diversity **(Figure 8D)**, indicating a selective recruitment of specific B-cell clones.

**Figure 8.**
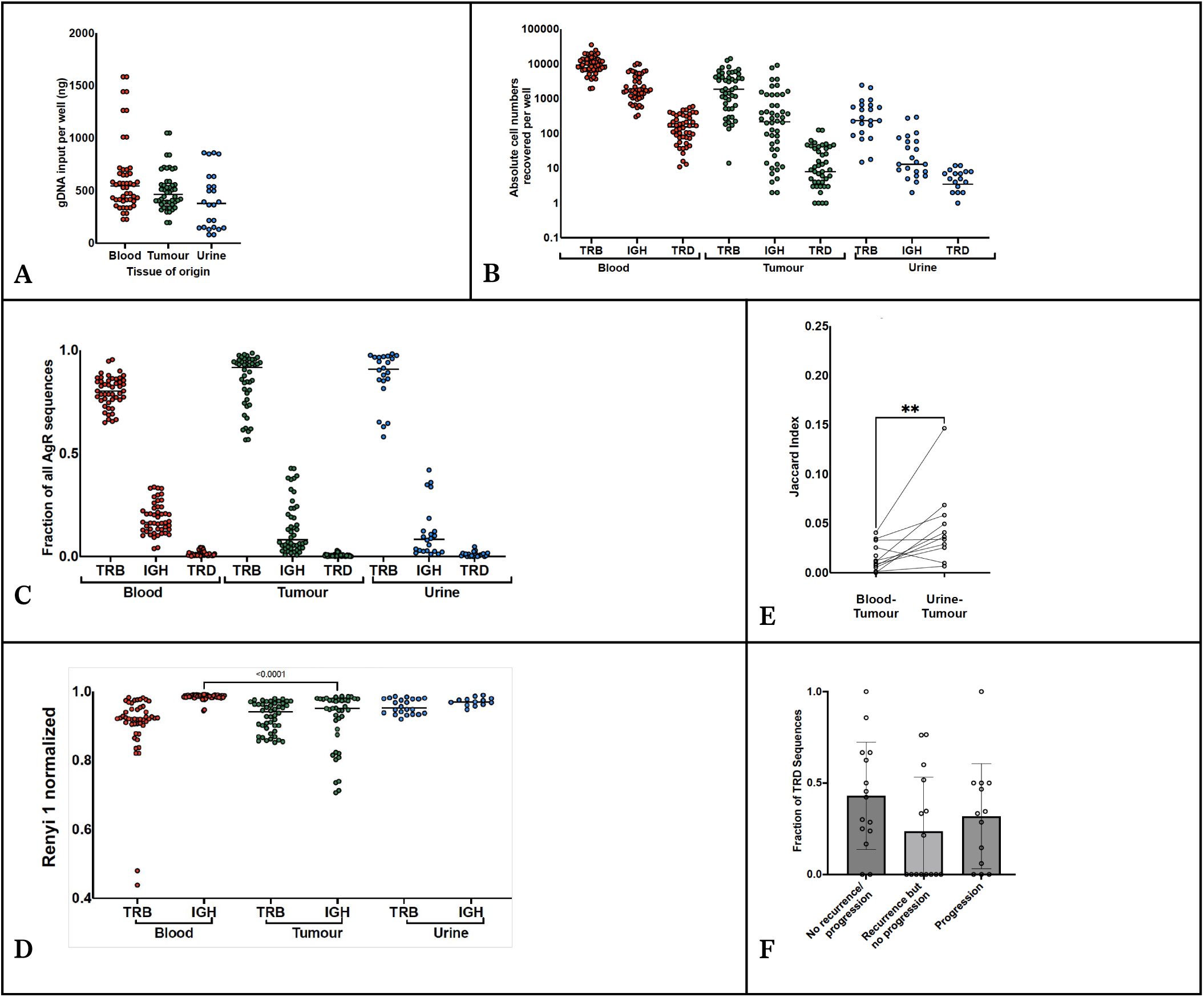
Immune profiles of blood, tumor and urine in NMIBC patients. **A.** Urine pellets (when available) yield less DNA compared with blood and tumor tissue. **B.** Cell recovery by lineage shows a similar trend, TRB > IGH > TRD, across all sample types with decreasing recovery across blood, tumor and urine as expected. **C.** Blood and tumor show distinct cell type representation: TRB is dominant in both, but at a higher average fraction in tumor tissue (AgR: Antigen Receptor). **D.** IGH repertoires in the tumor demonstrate significantly greater focusing than in blood. P-value from Kruskal-Wallis test on Renyi 1 normalized index with post-hoc Dunn’s test for cross tissue same chain comparisons. Only p-values less than 0.05 are shown. **E.** Tumor immune profile is more similar to urine than blood in samples with all sources available. **F.** Fraction of TRDV1 of total TRD sequences in different outcome cohorts shows a trend towards lower representation in recurrence or progression cohorts.

### 4.2. Representation of tumor immune repertoire in urine

Intriguingly, in the subset of patients with all three sample types available, the relative population frequencies and repertoire diversities in urine and tumor samples were more similar than tumor-blood comparisons. The clonal composition from urine samples more accurately reflected the clonal makeup of the tumor than the peripheral blood, with a greater degree of shared sequences and higher Jaccard Index for tumor-urine pairings compared to blood-tumor pairings **(Figure 8E)**. This suggests that immune cells found in the urine may be shed directly from the tumor tissue into the bladder, making urine a valuable, non-invasive proxy for tumor-infiltrating immune cells. (Wong et al., 2018) A preliminary analysis also identified a trend linking a lower fraction of the TRDV1 gene and a higher risk of disease progression or recurrence, highlighting the potential of γδ T cells as prognostic biomarkers in bladder cancer with a similar association observed in breast cancer (Wu et al., 2019) **(Figure 8F)**. This suggests the power of the assay to not only detect rare cell populations, but with a sensitivity that allows discrimination between patient samples and identification of potential prognostic associations.

### 4.3. Prognostic power of immune repertoire biomarkers

Despite the central role of lymphocytes in cancer surveillance, leveraging immune repertoire characteristics for clinical diagnostics is a nascent field. Bladder cancer is in particular a highly immunogenic malignancy characterized by an elevated tumor mutational burden (TMB) and dense immune infiltration within the tumor microenvironment (Kjær et al., 2025). Recent studies highlight that the diversity and clonality of immune receptor repertoires in peripheral blood can serve as powerful, non-invasive prognostic biomarkers (Charles et al., 2020; Valpione et al., 2021; Yang et al., 2025).

To assess the prognostic value of the immunoPETE data, we performed an analysis of progression-free survival using Cox regression. Due to the small cohort size, the analysis focused on immune biomarkers derived exclusively from blood samples to maximize the number of subjects included.

We compared three distinct models: Model 1 included only clinical factors (age, number of tumors, and size of the largest tumor); Model 2 included principal components (PCs) derived from 29 quantitative immune repertoire metrics (e.g., cell count, cell type ratios, and global and chain-specific diversity measures), selected to capture 90% of the total variance; and, Model 3 incorporated both the clinical factors from Model 1 and the PCs from Model 2. We compared the predictive power of these models using their concordance scores. Concordance score is a goodness-of-fit measure for time-to-event models and quantifies the fraction of sample pairs in which the model predicts the relative order of the event’s occurrence (cancer progression in this analysis) between the two samples correctly.

The concordance scores presented in **Table 3** suggest that immunoPETE biomarkers, even in this small cohort, may supplement traditional clinical factors in predicting patient outcomes. While these models currently rely on collective and quantitative metrics, future studies with larger cohorts could incorporate more specific data, such as the fraction of particular genes or the presence of specific clones, to enhance their predictive performance.

**Table 3.**
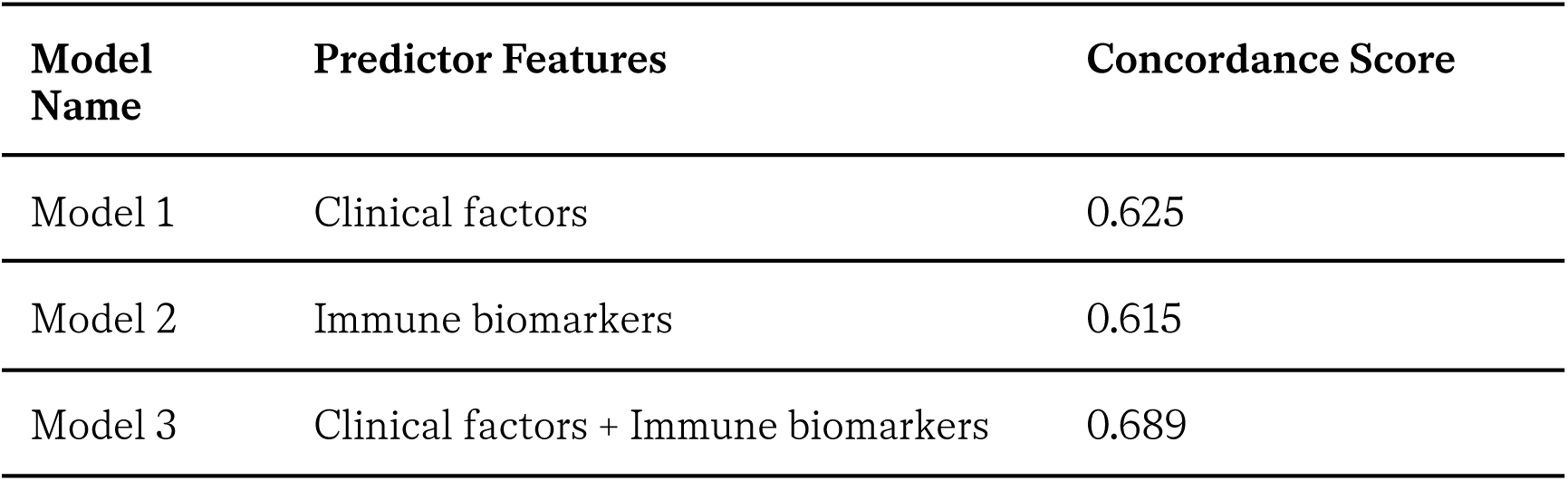
Predictive power of clinical factors and immune biomarkers towards patient outcome in the pilot NMIBC cohort.

| **Model Name** | **Predictor Features** | **Concordance Score** |
| --- | --- | --- |
| Model 1 | Clinical factors | 0.625 |
| Model 2 | Immune biomarkers | 0.615 |
| Model 3 | Clinical factors + Immune biomarkers | 0.689 |

## 5. Discussion

The characterization of the Adaptive Immune Receptor Repertoire (AIRR) represents a frontier in precision medicine, offering a high-resolution window into the host’s past exposures, current pathological states, and future therapeutic potential. In this study, we introduced and validated immunoPETE, a genomic DNA (gDNA)-based assay designed to provide a comprehensive and quantitative view of this repertoire across all B cell and T cell lineages.

### 5.1. The critical role of comprehensive repertoire profiling

As the “molecular record” of the adaptive immune system, V(D)J rearrangements reflect the clonal expansion of lymphocytes in response to antigens (Nielsen et al., 2019). Traditional methods often focus on single chains—primarily TRB or IGH—yet our results highlight the necessity of a multi-chain approach. By including the T-cell Receptor Delta (TRD) chain, immunoPETE captures γδ T cells, which occupy a unique niche at the intersection of innate and adaptive immunity. As seen in our NMIBC case study, the prognostic trend observed in TRDV1 usage underscores that focusing solely on αβ T cells may overlook critical effector populations in the tumor microenvironment (Chan, Duarte, Ostrouska, & Behren, 2022; Hayati et al., 2026).

### 5.2. Unique advantages of the immunoPETE architecture

A primary differentiator for immunoPETE is its reliance on gDNA rather than mRNA. While RNA-based assays offer high sensitivity for detecting rare transcripts, they are intrinsically biased by the transcriptional state of the cell; for instance, a single activated plasma cell can contain thousands of times more IGH mRNA than a resting B cell, leading to significant overestimation of clone sizes (Trück et al., 2021). By utilizing a 1:1 DNA-to-cell template ratio, immunoPETE provides a stable, internally normalized snapshot of the lymphocyte composition, including both active and bystander populations. This quantitative stability is further bolstered by the integration of Unique Molecular Identifiers (UMIs). Our data shows that UMIs effectively correct for PCR bias and sequencing errors, allowing for the accurate reconstruction of consensus sequences and the determination of absolute cell counts—prerequisites for identifying true biological variants and longitudinal monitoring of clonal evolution (Mazzotti et al., 2022).

The single-reaction integration of IGH, TRB, and TRD is a significant technical milestone. Most existing solutions require separate libraries for B and T cells, which complicates cross-comparison and consumes limited clinical specimens. The ability to observe B-T cell ratios and γδ T-cell dynamics within a single workflow provides a built-in control for sample quality and cellularity, ensuring that the relative proportions of infiltrating lymphocytes are captured with high fidelity.

### 5.3. Benchmarking and analytical fidelity

The comprehensive benchmarking reported here represents one of the most rigorous analytical validations of an AIRR-seq assay currently documented. While many studies focus on clinical cohorts, few provide the granularity offered here using multiple sample types (high complexity PBMC, low complexity HuT-78 cell line, contrived mixtures of PBMC and cell line, contrived mixtures of Pan-B and Pan-T samples) to investigate the accuracy and reproducibility of the assay in capturing key aspects of the immune repertoire, e.g. cell count, cell type ratio, gene usage and clonal composition.

Our findings—demonstrating strong linearity (R^2^ > 0.98) between DNA input and cell recovery across the 500–2500 ng range establish a clear operational window for clinical research (the “input amount” set). The linearity of the assay was shown to extend down to a 2 ng DNA input, as demonstrated by experiments utilizing a wider range of DNA amounts distributed on a logarithmic scale (the Limit of Quantitation, LoQ set). The spike-in experiments with HuT-78 cells further demonstrated a rare clone sensitivity of 10^-4^ (0.01%) using 2000 ng of input DNA (presumably reaching 10^-5^ using 20000 ng of input DNA). In the context of Minimal Residual Disease (MRD) monitoring, where detecting a single malignant clone among millions of healthy cells is paramount, this level of precision is essential (Pai & Satpathy, 2021; Seo & Choi, 2025). The reproducibility of top clone frequencies —even for clones representing less than 1% of the repertoire— confirms that immunoPETE is a reliable tool for longitudinal monitoring where stability of the baseline is critical for detecting meaningful immunological shifts (the “precision” set).

### 5.4. Clinical translation in oncology: Insights from bladder cancer

Our application of immunoPETE to a cohort of non-muscle invasive bladder cancer (NMIBC) patients highlights the value of multi-lineage profiling in solid tumors. We demonstrated that urine pellets can serve as a valuable, non-invasive “liquid biopsy” for the tumor-associated immune environment. The high Jaccard similarity observed between tumor and urine repertoires suggests that immune cells in the urine are directly shed from the bladder malignancy rather than reflecting systemic circulation.

The identification of a prognostic trend in TRDV1 gene usage among γδ T cells further underscores the importance of capturing “marginal” T-cell types. Often overlooked in traditional αβ-focused studies, γδ T cells are critical effectors in tumor surveillance (Hu et al., 2023). Moreover, integrating immune biomarkers with clinical factors improved the predictive concordance score from 0.625 to 0.689. This suggests that while clinical staging remains vital, the “immunological state”—defined by metrics such as TRB dominance or B-cell clonal focusing—provides independent, additive value in predicting progression-free survival. This aligns with recent findings in melanoma and lung cancer, where pre-treatment TCR diversity has been linked to better responses to immune checkpoint inhibitors (Valpione et al., 2021).

### 5.5. Broader applications and future directions

AIRR-seq has been transformative in resolving how the immune system adapts—or fails to adapt—to pathogens. During the COVID-19 pandemic, immunoPETE was instrumental in identifying immune factors associated with disease severity (Dannebaum et al., 2022) and showing that while B-cell narrowing is a predictable response to infection, a profound contraction of the T-cell repertoire in patients over 50 served as a major risk factor for severe disease (Joseph et al., 2022).

In autoimmunity, immunoPETE has clarified the pathophysiology of multiple sclerosis (MS). MS patients were found to harbor a uniquely broad EBV-specific TCR repertoire, supporting the hypothesis of dysregulated antiviral immunity as a disease driver (Schneider-Hohendorf et al., 2022). Furthermore, the assay’s ability to track B-cell reconstitution following anti-CD20 therapy (ocrelizumab) revealed that the repopulation of “memory” high-SHM clones predicted ongoing disease activity (Schneider-Hohendorf et al., 2025). This demonstrates how AIRR-seq can serve as a functional biomarker to monitor therapeutic efficacy and forecast disease progression on an individualized basis.

The scalability of immunoPETE was exemplified by the Human Blood Immunome Encyclopedia (HuBIE) (Akbari et al., 2025). Comprising 2,614 profiles, HuBIE provides an unprecedented baseline of real-world immune diversity. Large-scale datasets are essential for building population-level models of immune aging and identifying shared “public” clonotypes across diverse patient demographics. This resource fills a critical gap, offering the reference data needed to train machine-learning classifiers for future diagnostic and prognostic use.

The versatility of immune repertoire profiling extends into several other critical areas of precision medicine. By analyzing somatic hypermutation (SHM) and lineage trees, researchers can identify high-affinity neutralizing antibodies and quantify the duration of vaccine-induced memory (Nielsen et al., 2019). Repertoire features can serve as inclusion/exclusion criteria in clinical trials, e.g. identifying patients with the necessary TCR diversity to respond to immunotherapies like checkpoint inhibitors (Liu et al., 2024; Y. Li et al., 2021). Monitoring for specific clonal expansions has been shown to provide early warnings of organ rejection or graft-versus-host disease (GvHD) (DeWolf et al., 2018; Leick et al., 2020; Tian, Li, & Lv, 2021).

As sequencing costs continue to decrease while analytical frameworks mature, AIRR-seq is poised to become a routinely deployed clinical tool. Future directions include integrating bulk DNA profiling with single-cell transcriptomics and spatial biology to link clonal identity with functional phenotypes. Ultimately, tools like immunoPETE provide a unifying lens through which to understand how diverse disease processes perturb adaptive immunity, offering the critical insights needed to improve patient outcomes through precision immunology.

## 8. Supplementary Material

### Tables

**Table S1** Details of immunoPETE reports

**Table S2** Experimental design of the immunoPETE analytical performance characterization study

**Table S3** Cell type ratio measurements in Pan-T / Pan-B set

**Table S4** Sample summary statistics of PBMC1 and PBMC2

**Table S5** Identity and frequency of top 10 clones in PBMC1-4

**Table S6** Chi-squared test results for V and J gene pairing independence

**Table S7** Demographic and clinical description of the NMIBC cohort in the case study

**Table S8** Hill numbers of all replicates of PBMC1-4

### Figures

**Figure S1.**
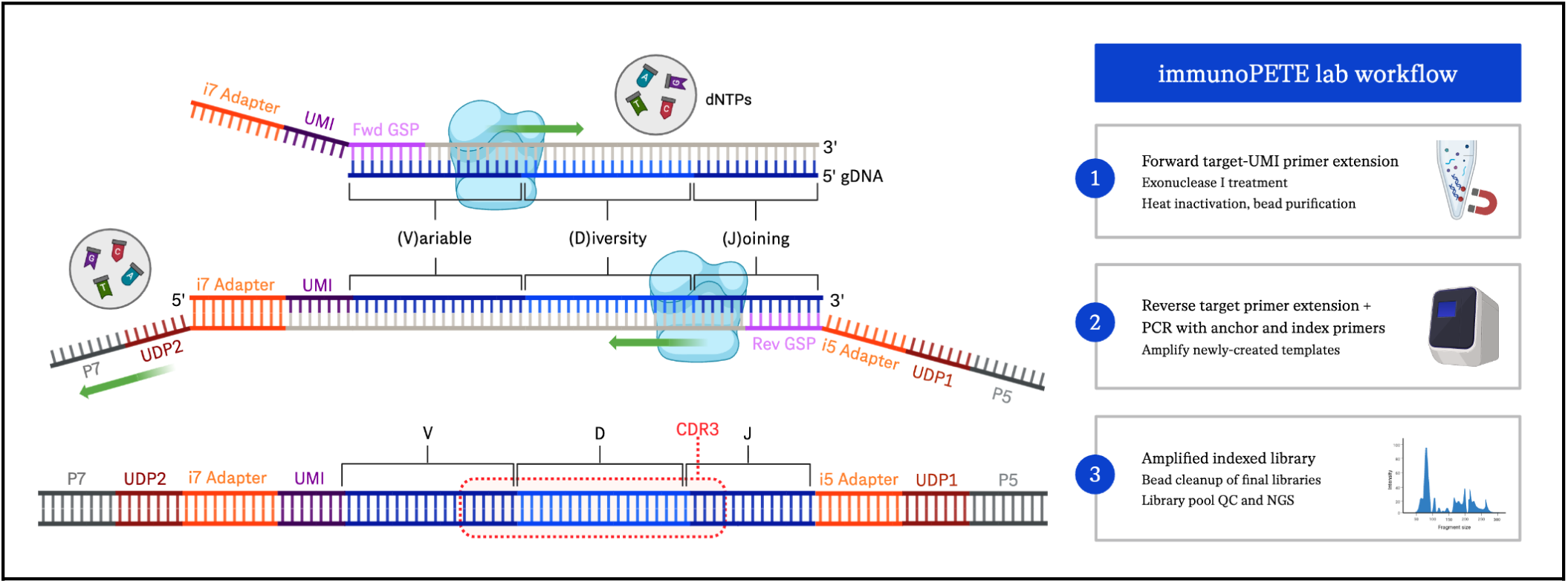
Primer extension and target enrichment workflow. immunoPETE uses a two-stage PCR process. First, V-gene oligos with unique identifiers extend the DNA, which is then purified, followed by a second extension with J-gene and indexed sequencing primers, and the resulting libraries are purified and pooled for sequencing.

**Figure S2.**
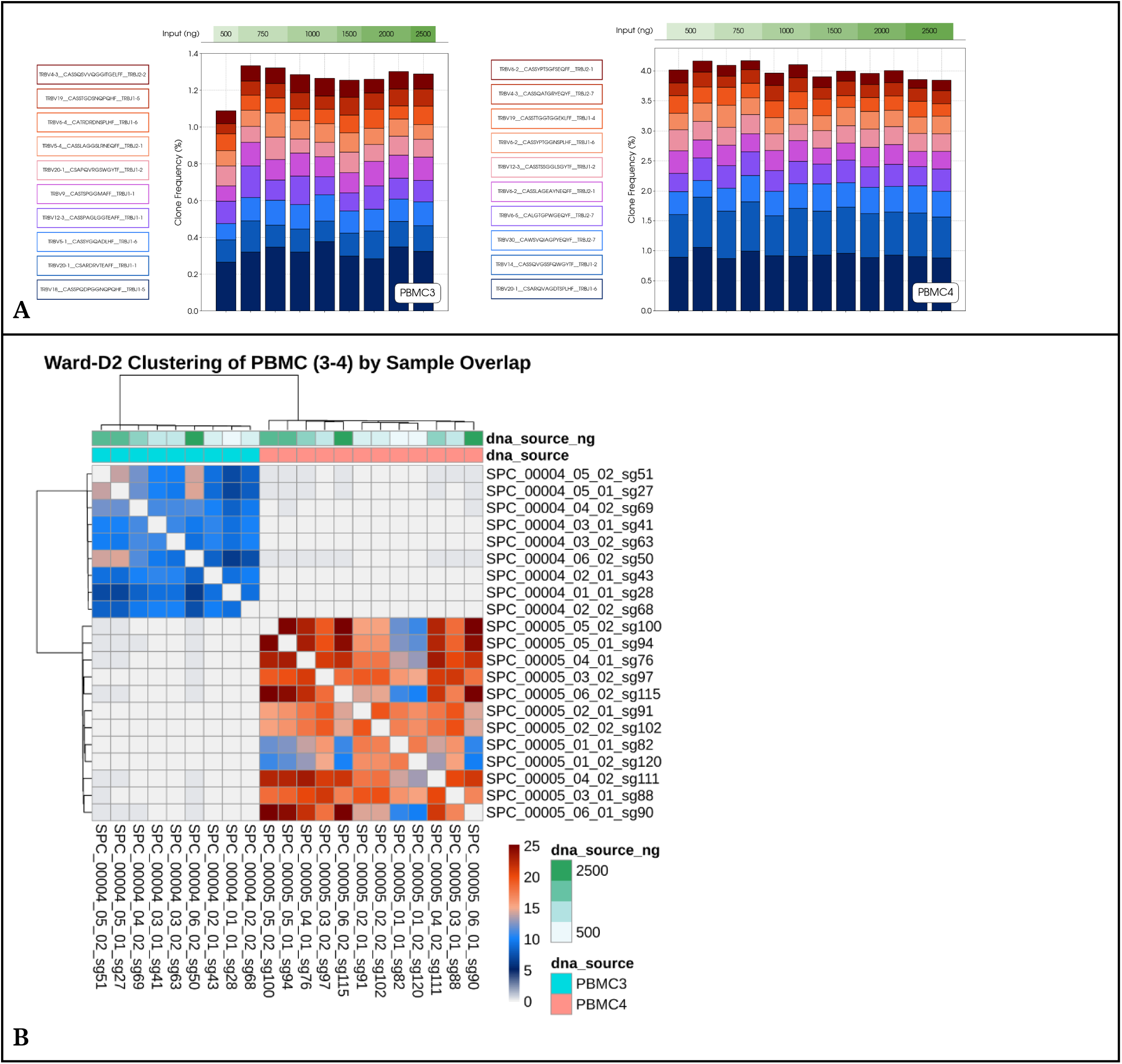
Analysis of clonal composition in High-Complexity (PBMC3-4) sample sets. **A.** Abundance of top 10 clones in PBMC3 and PBMC4 from 500 ng to 2500 ng of starting gDNA. The relative clone frequencies were consistent among replicates even for clones which comprise a smaller fraction of the overall repertoire, demonstrating reliable quantification of both dominant and rare clonotypes within the standard working range **(Table S5)**. **B.** Pairwise similarity for PBMC3 and PBMC4 libraries was represented by the Sorensen similarity index (Equation 2), characterized by the fraction of aggregated shared cell counts between two samples belonging to the same clonotype. Generally, higher overlap is observed in more dominant repertoires. See Table S2 for information on specific replicate description and attributes.

**Figure S3.**
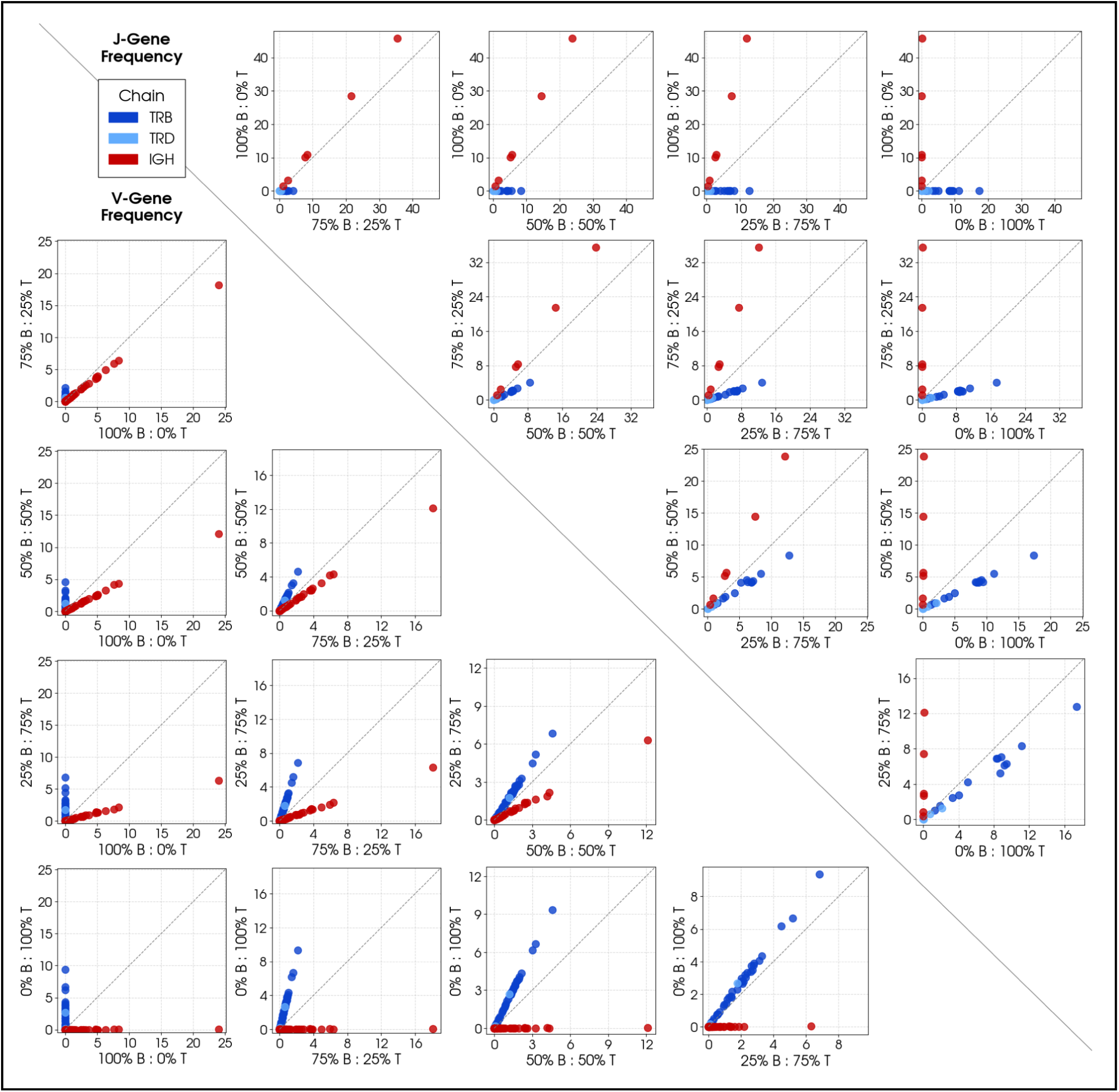
V/J-gene usage comparison in Pan-B and Pan-T mixtures. Percent frequency of all observed V-genes (bottom-left) and J-genes (upper-right) across five tested contrived mixtures of Pan-T and Pan-B cells. Each subplot illustrates pairwise gene frequencies, with points nearing the diagonal indicating similarity in abundance between the two mixtures. The distribution of IGH (red) points dominates the B-cell rich comparisons, while TR (blue) points become increasingly dominant as the T-cell fraction increases. We opted to combine and normalize gene frequencies within a single pool, encompassing IGH, TRB, and TRD classes, instead of the typical normalization within each class. This approach allows us to simultaneously represent cell type fractions and gene frequencies in a single plot for each comparison.

**Figure S4.**
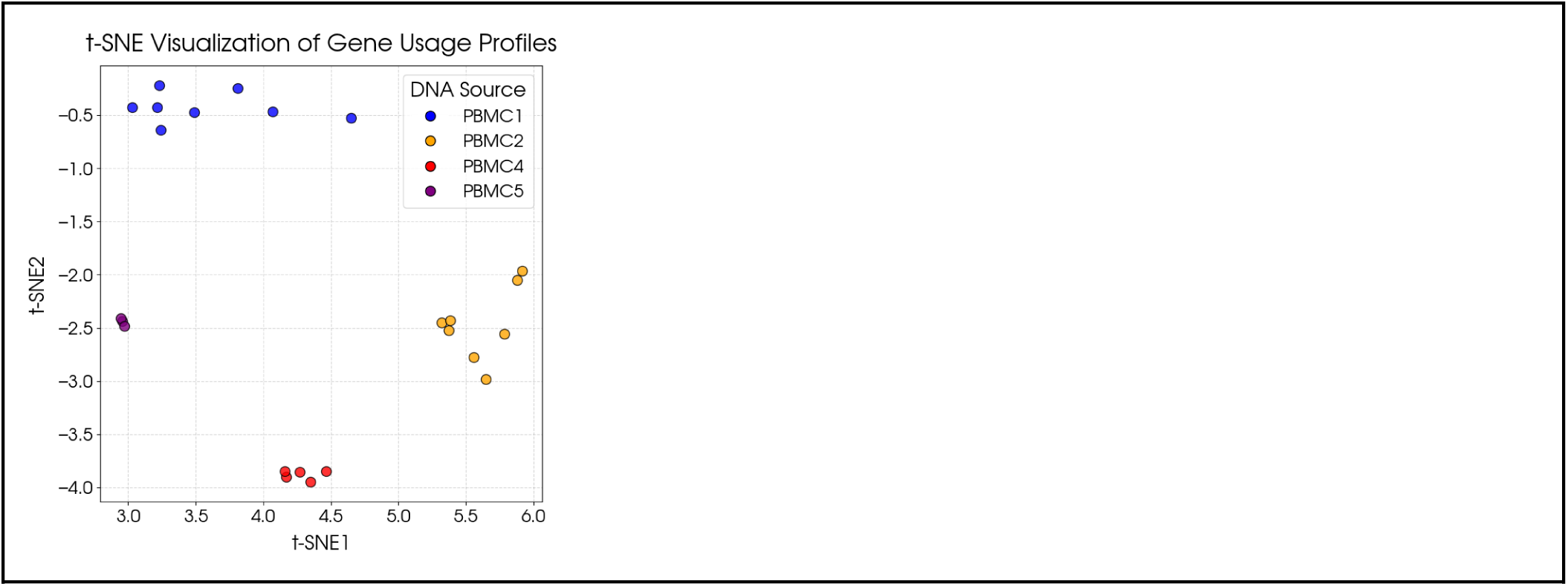
t-SNE projection of gene usage profiles. t-SNE projection of z-transformed gene usage across all 2000-ng PBMC replicates with a minimum of 3 libraries (PBMC1-2, 4-5). Clustering reveals distinct patterns in V/J gene expression among all samples, highlighting sample-specific variations in gene usage.

**Figure S5.**
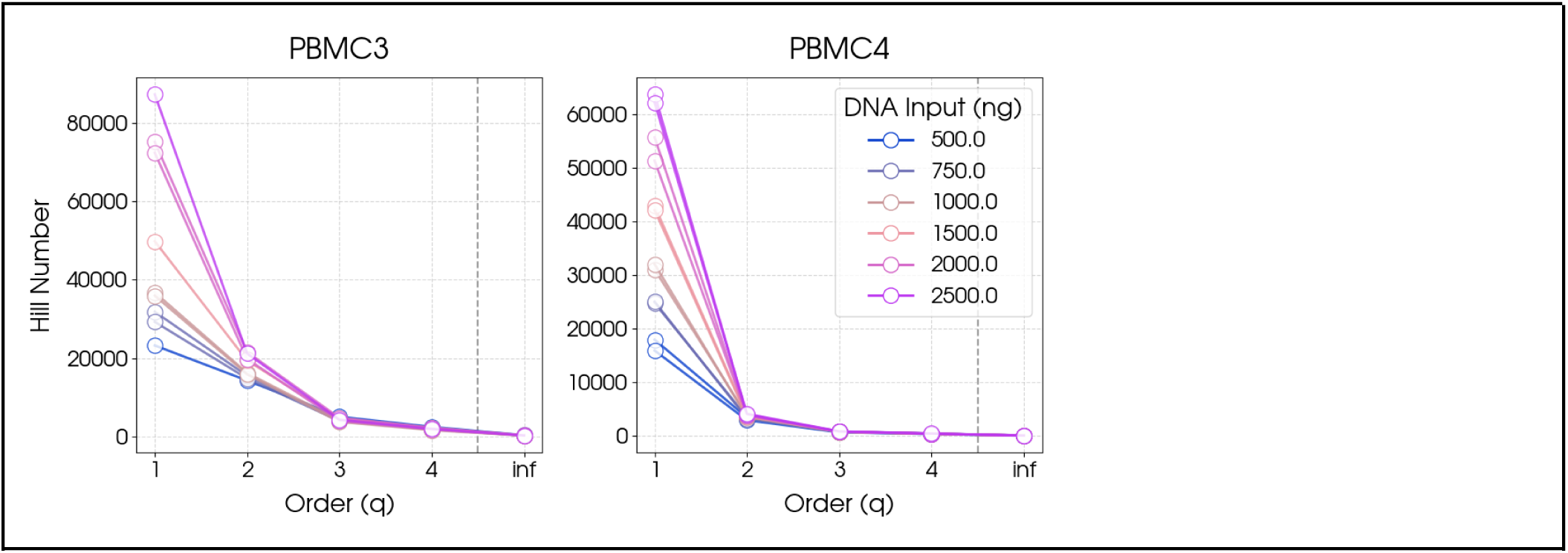
Hill numbers of all technical replicates in PBMC3 and PBMC4. TCR and BCR repertoire diversity quantified using Hill numbers (Effective Number of Clonotypes), where increasing values of q reflects a heavier weighting on dominant clonotypes. Diversity is measured at a range of q values from q=0 (Clonal richness), q=1 (Exponential of Shannon Index), to q=inf (inverse of the proportion of the most dominant clone). Full data is available in Table S8.

## Footnotes

1 The results of this study were presented as a poster in the Immune Responses in Cancer and Infection (IRCI) conference in 2022.

2 The samples and associated data were obtained through a material transfer agreement from the University of Birmingham’s Bladder Cancer Prognosis Programme, under the terms of the original ethical approval (MREC 06/MRE04/65).

