## Supplementary material for "immunoPETE: A DNA-based integrated B-cell and T-cell receptor profiling platform": Table S1

### Table S1 - Details of immunoPETE reports.

**Report Type: Sample Summary Report**

| PROPERTY_NAME | DESCRIPTION | EXAMPLE |
|---|---|---|
| experiment_name | The name assigned to the project (structure) | iPETE_LoDExp1_LibPool_SampleSheet_NextSeqHO_Run_081922 |
| sample_name | The unique identifier for each technical replicate (structure) | SPC_00008_01_01_sg8 |
| status | If the pipeline was completed for this replicate. In the case of a fail, the reason is provided.<br>(values: "pass", "fail: <10 cells", "fail: <50k reads") | pass |
| valid_on_target_reads | Total number of on-target reads with valid UMIs assigned to the sample replicate | 27660804 |
| UMI_families | Number of reported families after the deduplication step. A family is comprised of one or more valid on-target reads collapsed into a unique Vgene/CDR3/Jgene/UMI construct | 347373 |
| percent_functional | Percentage of UMI families that have a functional (productive) Vgene/CDR3/Jgene rearrangement | 80.8005092 |
| cell_count | Number of reported cells | 164986 |
| TRB | Number of reported TRB cells | 159166 |
| TRD | Number of reported TRD cells | 5438 |
| IGH | Number of reported IGH cells | 382 |
| chimera | Number of rearrangements where the Vgene and Jgene parts of the read align to regions in different chain types | 45 |
| IGH_percent | Fraction of cells classified as IGH (IGH_percent = IGH / cell_count) | 0.00231472 |
| TRB_percent | Fraction of cells classified as TRB (TRB_percent = TRB / cell_count) | 0.96446122 |
| TRD_percent | Fraction of cells classified as TRD (TRD_percent = TRD / cell_count) | 0.03295138 |
| unique_CDR3 | Number of unique representitive CDR3s | 125275 |
| D50 | After sorting by counts, the fraction of total clones that make up 50% of the sample content | 0.36419815 |
| gini_index | A measure of unevenness of clone counts in the sample. (gini index ~ 1 suggests that the sample is dominated by one or few clone with most others having very low frequencies, whereas gini index ~ 0 suggests that the clones in the sample are close to being evenly distributed) | 0.20446926 |
| renyi_1_norm | Renyi entropy of order 1 in the sample based on clone counts. A larger value suggests a higher degree of evenness. | 0.975362 |
| renyi_2_norm | Renyi entropy of order 2 in the sample based on clone counts. A larger value suggests a higher degree of evenness. | 0.865887 |
| simpsons_diversity_index | Measure of clone diversity within a sample. A larger value suggests a more diverse (evenly frequent) set of clones | 0.999969 |
| simpsons_dominance_index | Measure of clone dominance within a sample. A larger value suggests a few clonal types occupying a large proportion of all clones | 0.000037 |
| TRB_renyi_1_norm | renyi_1_norm for TRB cell type | 0.97873371 |
| TRB_renyi_2_norm | renyi_2_norm for TRB cell type | 0.88281642 |
| TRB_simpsons_diversity_norm | simpsons_diversity_norm for TRB cell type | 0.99997523 |
| TRB_simpsons_dominance_norm | simpsons_dominance_norm for TRB cell type | 3.11E-05 |
| TRB_unique_cdr3 | unique_cdr3 for TRB cell type | 127727 |
| TRB_D50 | D50 for TRB cell type | 0.37692892 |
| TRB_gini_index | gini_index for TRB cell type | 0.1889249 |
| IGH_renyi_1_norm | renyi_1_norm for IGH cell type | 0.99369129 |
| IGH_renyi_2_norm | renyi_2_norm for IGH cell type | 0.98348102 |
| IGH_simpsons_diversity_norm | simpsons_diversity_norm for IGH cell type | 0.99947781 |
| IGH_simpsons_dominance_norm | simpsons_dominance_norm for IGH cell type | 0.00313862 |
| IGH_unique_cdr3 | unique_cdr3 for IGH cell type | 351 |
| IGH_D50 | D50 for IGH cell type | 0.45584046 |
| IGH_gini_index | gini_index for IGH cell type | 0.07629659 |
| TRD_renyi_1_norm | renyi_1_norm for TRD cell type | 0.84616555 |
| TRD_renyi_2_norm | renyi_2_norm for TRD cell type | 0.65456345 |
| TRD_simpsons_diversity_norm | simpsons_diversity_norm for TRD cell type | 0.99238304 |
| TRD_simpsons_dominance_norm | simpsons_dominance_norm for TRD cell type | 0.00779945 |
| TRD_unique_cdr3 | unique_cdr3 for TRD cell type | 1661 |
| TRD_D50 | D50 for TRD cell type | 0.05478627 |
| TRD_gini_index | gini_index for TRD cell type | 0.62377335 |

| | | |
|---|---|---|
| **Report Type: Cell-level CDR3 Report** | | |
| PROPERTY_NAME | DESCRIPTION | EXAMPLE |
| biosample_name | Name of the sample specimen | PTL_12345_01 |
| sample_name | Name of the technical replicate the rearrangement belongs to | PTL_12345_01_02 |
| chain | Gene locus that codes for one of the protein chains of an adaptive immune receptor | IGH |
| n_reads | Number of reads (UMI family size) that were deduplicated to create the rearrangement | 521 |
| cdr3_nt | Representative CDR3 sequence of the rearrangement (nucleotide) | TGTGCGAGAGGCGACGATTTTTGGAGGGCGAGGTACTACTACGGTATGGACGTCTGG |
| cdr3_aa | Representative CDR3 sequence of the rearrangement (amino acid) | CARGDDFWRARYYYGMDVW |
| cdr3_qual | A string of ASCII characters representing the Phred quality scores for each base in the CDR3 sequence | ]]]]]]]]]]]]]]]]]]]]]]]]]]]]]]]]]]]]]]]]]]]]]]]]]]]]]]]]] |
| v_gene | Annotated V-gene | IGHV1-69D |
| multi_v | Number of unique alignments to the annotated V-gene | 3 |
| v_gene_class | Annotated V-gene product class | immunoglobulin gene |
| v_gene_type | Type of the V-gene (Values: gene, pseudogene, non-functional, putative-CDR3-boundary) | gene |
| d_gene | Annotated D-gene | |
| j_gene | Annotated J-gene | IGHJ6 |
| multi_j | Number of unique alignments to the annotated J-gene | 1 |
| j_gene_class | Annotated J-gene product class | immunoglobulin gene |
| j_gene_type | Type of the J-gene (Values: gene, pseudogene, non-functional, putative-CDR3-boundary) | gene |
| umi_seq | UMI sequence | TCACATAGCAA |
| consensus_seq | Nucleic acid sequence derived from an alignment of multiple individual reads | CACGAGCACAGCCTACATGGAGCTGAGCAGCCTGAGATCTGAGGACACGGCCGTGTATTACTGTGCGAGAGGCGACGATTTTTGGAGGGCGAGGTACTACTACGGTATGGACGTCTGGGGCCAAGGGACCACGGTCA |
| consensus_qual | A string of ASCII characters representing the Phred quality scores for each base in the consensus_seq | ]]]]]]]]]]]]]]]]]]]]]]]]]]]]]]]]]]]]]]]]]]]]]]]]]]]]]]]]]]]]]]]]]]]]]]]]]]]]]]]]]]]]]]]]]]]]]]]]]]]]]]]]]]]]]]]]]]]]]]]]]]]]]]]]]]]]]]]]] |
| consensus_loc | Location of the start of the consensus on the read | 62 |
| is_functional | Indication if the CDR3 sequence is a presumably productive rearrangement. | TRUE |
| cdr3_type | If the rearrangement is not productive, the reason(s) for non-productivity are listed, such as a stop codon or frameshift, separated by ";" | stop_codon;frameshift |
| clone_id | Unique identifier for the clone (e.g. Vgene__CDR3AA__Jgene) | IGHV1-69D__CARGDDFWRARYYYGMDVW__IGHJ6 |

| | | |
|---|---|---|
| **Report Type: Clone-level CDR3 Report** | | |
| PROPERTY_NAME | DESCRIPTION | EXAMPLE |
| biosample_name | Name of the sample specimen | PTL_12345_01 |
| sample_name | Name of the replicate | PTL_12345_01_02; PTL_12345_01_03 |
| chain | Gene locus that codes for one of the protein chains of an adaptive immune receptor | IGH |
| count | Number of cells or templates in the sample with this clonotype | 16 |
| n_reads | Number of reads (UMI family size) that were deduplicated to create the rearrangement | 521 |
| cdr3_nt | Representative CDR3 sequence of the rearrangement (nucleotide) | TGTGCGAGAGGCGACGATTTTTGGAGGGCGAGGTACTACTACGGTATGGACGTCTGG |
| cdr3_aa | Representative CDR3 sequence of the rearrangement (amino acid) | CARGDDFWRARYYYGMDVW |
| v_gene | Annotated V-gene | IGHV1-69D |
| v_gene_class | Annotated V-gene product class | immunoglobulin gene |
| v_gene_type | Type of the V-gene (Values: gene, pseudogene, non-functional, putative-CDR3-boundary) | gene |
| d_gene | Annotated D-gene | |
| j_gene | Annotated J-gene | IGHJ6 |
| j_gene_class | Annotated J-gene product class | immunoglobulin gene |
| j_gene_type | Type of the J-gene (Values: gene, pseudogene, non-functional, putative-CDR3-boundary) | gene |
| is_functional | Indication if the CDR3 sequence is a presumably productive rearrangement. | FALSE |
| cdr3_type | If the rearrangement is not productive, the reason(s) for non-productivity are listed, such as a stop codon or frameshift, separated by ";" | stop_codon;frameshift |
| clone_id | Unique identifier for the clone (e.g. Vgene__CDR3AA__Jgene) | IGHV1-69D__CARGDDFWRARYYYGMDVW__IGHJ6 |
