## Supplementary material for "immunoPETE: A DNA-based integrated B-cell and T-cell receptor profiling platform": Table S2

**Table S2 - Experimental design of the immunoPETE analytical performance characterization study.**

| Replicate ID | Biosample ID | Subject ID | Sample processing batch | Position on plate | High complexity DNA source | High complexity DNA source type | High complexity input DNA amount (ng) | Low complexity DNA source | Low complexity DNA amount (ng) | Low complexity DNA fraction | Expected Hut78 cell count | Sample set "nickname" | Purpose |
|---|---|---|---|---|---|---|---|---|---|---|---|---|---|
| SPC_00000_01_01 | SPC_00000_01 | SPC_00000 | 1 | C5 | None | None | 0 | None | 0 | 0 | 0 | NTC | Detect contamination; LoB |
| SPC_00000_01_02 | SPC_00000_01 | SPC_00000 | 1 | C6 | None | None | 0 | None | 0 | 0 | 0 | NTC | Detect contamination; LoB |
| SPC_00000_01_03 | SPC_00000_01 | SPC_00000 | 2 | C7 | None | None | 0 | None | 0 | 0 | 0 | NTC | Detect contamination; LoB |
| SPC_00000_01_04 | SPC_00000_01 | SPC_00000 | 3 | A1 | None | None | 0 | None | 0 | 0 | 0 | NTC | Detect contamination; LoB |
| SPC_00000_01_05 | SPC_00000_01 | SPC_00000 | 4 | C8 | None | None | 0 | None | 0 | 0 | 0 | NTC | Detect contamination; LoB |
| SPC_00000_01_06 | SPC_00000_01 | SPC_00000 | 4 | C5 | None | None | 0 | None | 0 | 0 | 0 | NTC | Detect contamination; LoB |
| SPC_00000_01_07 | SPC_00000_01 | SPC_00000 | 5 | B6 | None | None | 0 | None | 0 | 0 | 0 | NTC | Detect contamination; LoB |
| SPC_00000_01_08 | SPC_00000_01 | SPC_00000 | 5 | A7 | None | None | 0 | None | 0 | 0 | 0 | NTC | Detect contamination; LoB |
| SPC_00000_01_09 | SPC_00000_01 | SPC_00000 | 5 | B4 | None | None | 0 | None | 0 | 0 | 0 | NTC | Detect contamination; LoB |
| SPC_00001_01_01 | SPC_00001_01 | SPC_00001 | 1 | A6 | None | None | 0 | Hot78 cell line | 0.02 | 1 | 3 | Low Complexity | Low complexity sample: LoD; LoQ; linearity on log scale; precision; accuracy; contamination; UMI differentiating limit |
| SPC_00001_01_02 | SPC_00001_01 | SPC_00001 | 2 | B4 | None | None | 0 | Hot78 cell line | 0.02 | 1 | 3 | Low Complexity | Low complexity sample: LoD; LoQ; linearity on log scale; precision; accuracy; contamination; UMI differentiating limit |
| SPC_00001_01_03 | SPC_00001_01 | SPC_00001 | 4 | A3 | None | None | 0 | Hot78 cell line | 0.02 | 1 | 3 | Low Complexity | Low complexity sample: LoD; LoQ; linearity on log scale; precision; accuracy; contamination; UMI differentiating limit |
| SPC_00001_01_04 | SPC_00001_01 | SPC_00001 | 5 | B1 | None | None | 0 | Hot78 cell line | 0.02 | 1 | 3 | Low Complexity | Low complexity sample: LoD; LoQ; linearity on log scale; precision; accuracy; contamination; UMI differentiating limit |
| SPC_00001_02_01 | SPC_00001_02 | SPC_00001 | 1 | C8 | None | None | 0 | Hot78 cell line | 0.2 | 1 | 33 | Low Complexity | Low complexity sample: LoD; LoQ; linearity on log scale; precision; accuracy; contamination; UMI differentiating limit |
| SPC_00001_02_02 | SPC_00001_02 | SPC_00001 | 2 | B2 | None | None | 0 | Hot78 cell line | 0.2 | 1 | 33 | Low Complexity | Low complexity sample: LoD; LoQ; linearity on log scale; precision; accuracy; contamination; UMI differentiating limit |
| SPC_00001_02_03 | SPC_00001_02 | SPC_00001 | 4 | A1 | None | None | 0 | Hot78 cell line | 0.2 | 1 | 33 | Low Complexity | Low complexity sample: LoD; LoQ; linearity on log scale; precision; accuracy; contamination; UMI differentiating limit |
| SPC_00001_02_04 | SPC_00001_02 | SPC_00001 | 5 | C5 | None | None | 0 | Hot78 cell line | 0.2 | 1 | 33 | Low Complexity | Low complexity sample: LoD; LoQ; linearity on log scale; precision; accuracy; contamination; UMI differentiating limit |
| SPC_00001_03_01 | SPC_00001_03 | SPC_00001 | 1 | B3 | None | None | 0 | Hot78 cell line | 2 | 1 | 333 | Low Complexity | Low complexity sample: LoD; LoQ; linearity on log scale; precision; accuracy; contamination; UMI differentiating limit |
| SPC_00001_03_02 | SPC_00001_03 | SPC_00001 | 2 | A1 | None | None | 0 | Hot78 cell line | 2 | 1 | 333 | Low Complexity | Low complexity sample: LoD; LoQ; linearity on log scale; precision; accuracy; contamination; UMI differentiating limit |
| SPC_00001_03_03 | SPC_00001_03 | SPC_00001 | 4 | B5 | None | None | 0 | Hot78 cell line | 2 | 1 | 333 | Low Complexity | Low complexity sample: LoD; LoQ; linearity on log scale; precision; accuracy; contamination; UMI differentiating limit |
| SPC_00001_03_04 | SPC_00001_03 | SPC_00001 | 5 | B2 | None | None | 0 | Hot78 cell line | 2 | 1 | 333 | Low Complexity | Low complexity sample: LoD; LoQ; linearity on log scale; precision; accuracy; contamination; UMI differentiating limit |
| SPC_00001_04_01 | SPC_00001_04 | SPC_00001 | 1 | C1 | None | None | 0 | Hot78 cell line | 20 | 1 | 3333 | Low Complexity | Low complexity sample: LoD; LoQ; linearity on log scale; precision; accuracy; contamination; UMI differentiating limit |
| SPC_00001_04_02 | SPC_00001_04 | SPC_00001 | 2 | C2 | None | None | 0 | Hot78 cell line | 20 | 1 | 3333 | Low Complexity | Low complexity sample: LoD; LoQ; linearity on log scale; precision; accuracy; contamination; UMI differentiating limit |
| SPC_00001_04_03 | SPC_00001_04 | SPC_00001 | 4 | C1 | None | None | 0 | Hot78 cell line | 20 | 1 | 3333 | Low Complexity | Low complexity sample: LoD; LoQ; linearity on log scale; precision; accuracy; contamination; UMI differentiating limit |
| SPC_00001_04_04 | SPC_00001_04 | SPC_00001 | 5 | A8 | None | None | 0 | Hot78 cell line | 20 | 1 | 3333 | Low Complexity | Low complexity sample: LoD; LoQ; linearity on log scale; precision; accuracy; contamination; UMI differentiating limit |
| SPC_00001_05_01 | SPC_00001_05 | SPC_00001 | 1 | B4 | None | None | 0 | Hot78 cell line | 200 | 1 | 33333 | Low Complexity | Low complexity sample: LoD; LoQ; linearity on log scale; precision; accuracy; contamination; UMI differentiating limit |
| SPC_00001_05_02 | SPC_00001_05 | SPC_00001 | 2 | C8 | None | None | 0 | Hot78 cell line | 200 | 1 | 33333 | Low Complexity | Low complexity sample: LoD; LoQ; linearity on log scale; precision; accuracy; contamination; UMI differentiating limit |
| SPC_00001_05_03 | SPC_00001_05 | SPC_00001 | 4 | B3 | None | None | 0 | Hot78 cell line | 200 | 1 | 33333 | Low Complexity | Low complexity sample: LoD; LoQ; linearity on log scale; precision; accuracy; contamination; UMI differentiating limit |
| SPC_00001_05_04 | SPC_00001_05 | SPC_00001 | 5 | C7 | None | None | 0 | Hot78 cell line | 200 | 1 | 33333 | Low Complexity | Low complexity sample: LoD; LoQ; linearity on log scale; precision; accuracy; contamination; UMI differentiating limit |
| SPC_00002_01_01 | SPC_00002_01 | SPC_00002 | 4 | B6 | PBMC1 | PBMC | 2000 | None | 0 | 0 | 0 | Precision | Precision (reproducibility); recovery of rare clones |
| SPC_00002_01_02 | SPC_00002_01 | SPC_00002 | 5 | A3 | PBMC1 | PBMC | 2000 | None | 0 | 0 | 0 | Precision | Precision (reproducibility); recovery of rare clones |
| SPC_00002_02_01 | SPC_00002_02 | SPC_00002 | 4 | A2 | PBMC1 | PBMC | 2000 | Hot78 cell line | 0.02 | 1.00E-05 | 3 | Precision | Precision (reproducibility); recovery of rare clones |
| SPC_00002_02_02 | SPC_00002_02 | SPC_00002 | 5 | B8 | PBMC1 | PBMC | 2000 | Hot78 cell line | 0.02 | 1.00E-05 | 3 | Precision | Precision (reproducibility); recovery of rare clones |
| SPC_00002_03_01 | SPC_00002_03 | SPC_00002 | 4 | B1 | PBMC1 | PBMC | 2000 | Hot78 cell line | 0.2 | 1.00E-04 | 33 | Precision | Precision (reproducibility); recovery of rare clones |
| SPC_00002_03_02 | SPC_00002_03 | SPC_00002 | 5 | C6 | PBMC1 | PBMC | 2000 | Hot78 cell line | 0.2 | 1.00E-04 | 33 | Precision | Precision (reproducibility); recovery of rare clones |
| SPC_00002_04_01 | SPC_00002_04 | SPC_00002 | 4 | A7 | PBMC1 | PBMC | 2000 | Hot78 cell line | 2 | 1.00E-03 | 333 | Precision | Precision (reproducibility); recovery of rare clones |
| SPC_00002_04_02 | SPC_00002_04 | SPC_00002 | 5 | A2 | PBMC1 | PBMC | 2000 | Hot78 cell line | 2 | 1.00E-03 | 333 | Precision | Precision (reproducibility); recovery of rare clones |
| SPC_00003_01_01 | SPC_00003_01 | SPC_00003 | 4 | B7 | PBMC2 | PBMC | 2000 | None | 0 | 0 | 0 | Precision | Precision (reproducibility); recovery of rare clones |
| SPC_00003_01_02 | SPC_00003_01 | SPC_00003 | 5 | B5 | PBMC2 | PBMC | 2000 | None | 0 | 0 | 0 | Precision | Precision (reproducibility); recovery of rare clones |
| SPC_00003_02_01 | SPC_00003_02 | SPC_00003 | 4 | B4 | PBMC2 | PBMC | 2000 | Hot78 cell line | 0.02 | 1.00E-05 | 3 | Precision | Precision (reproducibility); recovery of rare clones |
| SPC_00003_02_02 | SPC_00003_02 | SPC_00003 | 5 | C4 | PBMC2 | PBMC | 2000 | Hot78 cell line | 0.02 | 1.00E-05 | 3 | Precision | Precision (reproducibility); recovery of rare clones |
| SPC_00003_03_01 | SPC_00003_03 | SPC_00003 | 4 | A6 | PBMC2 | PBMC | 2000 | Hot78 cell line | 0.2 | 1.00E-04 | 33 | Precision | Precision (reproducibility); recovery of rare clones |
| SPC_00003_03_02 | SPC_00003_03 | SPC_00003 | 5 | C1 | PBMC2 | PBMC | 2000 | Hot78 cell line | 0.2 | 1.00E-04 | 33 | Precision | Precision (reproducibility); recovery of rare clones |
| SPC_00003_04_01 | SPC_00003_04 | SPC_00003 | 4 | A5 | PBMC2 | PBMC | 2000 | Hot78 cell line | 2 | 1.00E-03 | 333 | Precision | Precision (reproducibility); recovery of rare clones |
| SPC_00003_04_02 | SPC_00003_04 | SPC_00003 | 5 | A5 | PBMC2 | PBMC | 2000 | Hot78 cell line | 2 | 1.00E-03 | 333 | Precision | Precision (reproducibility); recovery of rare clones |
| SPC_00004_01_01 | SPC_00004_01 | SPC_00004 | 2 | A4 | PBMC3 | PBMC | 500 | None | 0 | 0 | 0 | Input amount | Precision (reproducibility); yield estimation and linearity assessment within a practical range of input masses |
| SPC_00004_01_02 | SPC_00004_01 | SPC_00004 | 3 | C1 | PBMC3 | PBMC | 500 | None | 0 | 0 | 0 | Input amount | Precision (reproducibility); yield estimation and linearity assessment within a practical range of input masses |
| SPC_00004_02_01 | SPC_00004_02 | SPC_00004 | 2 | C3 | PBMC3 | PBMC | 750 | None | 0 | 0 | 0 | Input amount | Precision (reproducibility); yield estimation and linearity assessment within a practical range of input masses |
| SPC_00004_02_02 | SPC_00004_02 | SPC_00004 | 3 | C4 | PBMC3 | PBMC | 750 | None | 0 | 0 | 0 | Input amount | Precision (reproducibility); yield estimation and linearity assessment within a practical range of input masses |
| SPC_00004_03_01 | SPC_00004_03 | SPC_00004 | 2 | C1 | PBMC3 | PBMC | 1000 | None | 0 | 0 | 0 | Input amount | Precision (reproducibility); yield estimation and linearity assessment within a practical range of input masses |
| SPC_00004_03_02 | SPC_00004_03 | SPC_00004 | 3 | B7 | PBMC3 | PBMC | 1000 | None | 0 | 0 | 0 | Input amount | Precision (reproducibility); yield estimation and linearity assessment within a practical range of input masses |
| SPC_00004_04_01 | SPC_00004_04 | SPC_00004 | 2 | B8 | PBMC3 | PBMC | 1500 | None | 0 | 0 | 0 | Input amount | Precision (reproducibility); yield estimation and linearity assessment within a practical range of input masses |
| SPC_00004_04_02 | SPC_00004_04 | SPC_00004 | 3 | C5 | PBMC3 | PBMC | 1500 | None | 0 | 0 | 0 | Input amount | Precision (reproducibility); yield estimation and linearity assessment within a practical range of input masses |
| SPC_00004_05_01 | SPC_00004_05 | SPC_00004 | 2 | A3 | PBMC3 | PBMC | 2000 | None | 0 | 0 | 0 | Input amount | Precision (reproducibility); yield estimation and linearity assessment within a practical range of input masses |
| SPC_00004_05_02 | SPC_00004_05 | SPC_00004 | 3 | A3 | PBMC3 | PBMC | 2000 | None | 0 | 0 | 0 | Input amount | Precision (reproducibility); yield estimation and linearity assessment within a practical range of input masses |
| SPC_00004_06_01 | SPC_00004_06 | SPC_00004 | 2 | B5 | PBMC3 | PBMC | 2500 | None | 0 | 0 | 0 | Input amount | Precision (reproducibility); yield estimation and linearity assessment within a practical range of input masses |
| SPC_00004_06_02 | SPC_00004_06 | SPC_00004 | 3 | A2 | PBMC3 | PBMC | 2500 | None | 0 | 0 | 0 | Input amount | Precision (reproducibility); yield estimation and linearity assessment within a practical range of input masses |
| SPC_00005_01_01 | SPC_00005_01 | SPC_00005 | 4 | B2 | PBMC4 | PBMC | 500 | None | 0 | 0 | 0 | Input amount | Precision (reproducibility); yield estimation and linearity assessment within a practical range of input masses |
| SPC_00005_01_02 | SPC_00005_01 | SPC_00005 | 5 | C8 | PBMC4 | PBMC | 500 | None | 0 | 0 | 0 | Input amount | Precision (reproducibility); yield estimation and linearity assessment within a practical range of input masses |
| SPC_00005_02_01 | SPC_00005_02 | SPC_00005 | 4 | C3 | PBMC4 | PBMC | 750 | None | 0 | 0 | 0 | Input amount | Precision (reproducibility); yield estimation and linearity assessment within a practical range of input masses |
| SPC_00005_02_02 | SPC_00005_02 | SPC_00005 | 5 | A6 | PBMC4 | PBMC | 750 | None | 0 | 0 | 0 | Input amount | Precision (reproducibility); yield estimation and linearity assessment within a practical range of input masses |
| SPC_00005_03_01 | SPC_00005_03 | SPC_00005 | 4 | B8 | PBMC4 | PBMC | 1000 | None | 0 | 0 | 0 | Input amount | Precision (reproducibility); yield estimation and linearity assessment within a practical range of input masses |
| SPC_00005_03_02 | SPC_00005_03 | SPC_00005 | 5 | A1 | PBMC4 | PBMC | 1000 | None | 0 | 0 | 0 | Input amount | Precision (reproducibility); yield estimation and linearity assessment within a practical range of input masses |
| SPC_00005_04_01 | SPC_00005_04 | SPC_00005 | 4 | A4 | PBMC4 | PBMC | 1500 | None | 0 | 0 | 0 | Input amount | Precision (reproducibility); yield estimation and linearity assessment within a practical range of input masses |
| SPC_00005_04_02 | SPC_00005_04 | SPC_00005 | 5 | B7 | PBMC4 | PBMC | 1500 | None | 0 | 0 | 0 | Input amount | Precision (reproducibility); yield estimation and linearity assessment within a practical range of input masses |
| SPC_00005_05_01 | SPC_00005_05 | SPC_00005 | 4 | C6 | PBMC4 | PBMC | 2000 | None | 0 | 0 | 0 | Input amount | Precision (reproducibility); yield estimation and linearity assessment within a practical range of input masses |
| SPC_00005_05_02 | SPC_00005_05 | SPC_00005 | 5 | A4 | PBMC4 | PBMC | 2000 | None | 0 | 0 | 0 | Input amount | Precision (reproducibility); yield estimation and linearity assessment within a practical range of input masses |
| SPC_00005_06_01 | SPC_00005_06 | SPC_00005 | 4 | C2 | PBMC4 | PBMC | 2500 | None | 0 | 0 | 0 | Input amount | Precision (reproducibility); yield estimation and linearity assessment within a practical range of input masses |
| SPC_00005_06_02 | SPC_00005_06 | SPC_00005 | 5 | C3 | PBMC4 | PBMC | 2500 | None | 0 | 0 | 0 | Input amount | Precision (reproducibility); yield estimation and linearity assessment within a practical range of input masses |
| SPC_00005_07_01 | SPC_00005_07 | SPC_00005 | 1 | A3 | PBMC4 | PBMC | 0.02 | Hot78 cell line | 2 | 0.9900990099 | 333 | Limit of Quantitation | High complexity samples: Precision (reproducibility); linearity on log scale; fraction of spiked-in Hut78 cells vs PBMC; LoD; LoQ |

**Table S2 - Experimental design of the immunoPETE analytical performance characterization study.**

| Replicate ID | Biosample ID | Subject ID | Sample processing batch | Position on plate | High complexity DNA source | High complexity DNA source type | High complexity input DNA amount (ng) | Low complexity DNA source | Low complexity DNA amount (ng) | Low complexity DNA fraction | Expected Hut78 cell count | Sample set "nickname" | Purpose |
|---|---|---|---|---|---|---|---|---|---|---|---|---|---|
| SPC_00005_07_02 | SPC_00005_07 | SPC_00005 | 2 | A6 | PBMC4 | PBMC | 0.02 | Hot78 cell line | 2 | 0.9900990099 | 333 | Limit of Quantitation | High complexity samples: Precision (reproducibility); linearity on log scale; fraction of spiked-in Hut78 cells vs PBMC; LoD; LoQ |
| SPC_00005_07_03 | SPC_00005_07 | SPC_00005 | 3 | B2 | PBMC4 | PBMC | 0.02 | Hot78 cell line | 2 | 0.9900990099 | 333 | Limit of Quantitation | High complexity samples: Precision (reproducibility); linearity on log scale; fraction of spiked-in Hut78 cells vs PBMC; LoD; LoQ |
| SPC_00005_08_01 | SPC_00005_08 | SPC_00005 | 1 | B1 | PBMC4 | PBMC | 0.2 | Hot78 cell line | 2 | 0.9090909091 | 333 | Limit of Quantitation | High complexity samples: Precision (reproducibility); linearity on log scale; fraction of spiked-in Hut78 cells vs PBMC; LoD; LoQ |
| SPC_00005_08_02 | SPC_00005_08 | SPC_00005 | 2 | B7 | PBMC4 | PBMC | 0.2 | Hot78 cell line | 2 | 0.9090909091 | 333 | Limit of Quantitation | High complexity samples: Precision (reproducibility); linearity on log scale; fraction of spiked-in Hut78 cells vs PBMC; LoD; LoQ |
| SPC_00005_08_03 | SPC_00005_08 | SPC_00005 | 3 | B4 | PBMC4 | PBMC | 0.2 | Hot78 cell line | 2 | 0.9090909091 | 333 | Limit of Quantitation | High complexity samples: Precision (reproducibility); linearity on log scale; fraction of spiked-in Hut78 cells vs PBMC; LoD; LoQ |
| SPC_00005_09_01 | SPC_00005_09 | SPC_00005 | 1 | A2 | PBMC4 | PBMC | 2 | Hot78 cell line | 2 | 0.5 | 333 | Limit of Quantitation | High complexity samples: Precision (reproducibility); linearity on log scale; fraction of spiked-in Hut78 cells vs PBMC; LoD; LoQ |
| SPC_00005_09_02 | SPC_00005_09 | SPC_00005 | 2 | A8 | PBMC4 | PBMC | 2 | Hot78 cell line | 2 | 0.5 | 333 | Limit of Quantitation | High complexity samples: Precision (reproducibility); linearity on log scale; fraction of spiked-in Hut78 cells vs PBMC; LoD; LoQ |
| SPC_00005_09_03 | SPC_00005_09 | SPC_00005 | 3 | C3 | PBMC4 | PBMC | 2 | Hot78 cell line | 2 | 0.5 | 333 | Limit of Quantitation | High complexity samples: Precision (reproducibility); linearity on log scale; fraction of spiked-in Hut78 cells vs PBMC; LoD; LoQ |
| SPC_00005_10_01 | SPC_00005_10 | SPC_00005 | 1 | B8 | PBMC4 | PBMC | 20 | Hot78 cell line | 2 | 0.09090909091 | 333 | Limit of Quantitation | High complexity samples: Precision (reproducibility); linearity on log scale; fraction of spiked-in Hut78 cells vs PBMC; LoD; LoQ |
| SPC_00005_10_02 | SPC_00005_10 | SPC_00005 | 2 | B1 | PBMC4 | PBMC | 20 | Hot78 cell line | 2 | 0.09090909091 | 333 | Limit of Quantitation | High complexity samples: Precision (reproducibility); linearity on log scale; fraction of spiked-in Hut78 cells vs PBMC; LoD; LoQ |
| SPC_00005_10_03 | SPC_00005_10 | SPC_00005 | 3 | A8 | PBMC4 | PBMC | 20 | Hot78 cell line | 2 | 0.09090909091 | 333 | Limit of Quantitation | High complexity samples: Precision (reproducibility); linearity on log scale; fraction of spiked-in Hut78 cells vs PBMC; LoD; LoQ |
| SPC_00005_11_01 | SPC_00005_11 | SPC_00005 | 1 | B5 | PBMC4 | PBMC | 200 | Hot78 cell line | 2 | 0.009900990099 | 333 | Limit of Quantitation | High complexity samples: Precision (reproducibility); linearity on log scale; fraction of spiked-in Hut78 cells vs PBMC; LoD; LoQ |
| SPC_00005_11_02 | SPC_00005_11 | SPC_00005 | 2 | A2 | PBMC4 | PBMC | 200 | Hot78 cell line | 2 | 0.009900990099 | 333 | Limit of Quantitation | High complexity samples: Precision (reproducibility); linearity on log scale; fraction of spiked-in Hut78 cells vs PBMC; LoD; LoQ |
| SPC_00005_11_03 | SPC_00005_11 | SPC_00005 | 3 | C2 | PBMC4 | PBMC | 200 | Hot78 cell line | 2 | 0.009900990099 | 333 | Limit of Quantitation | High complexity samples: Precision (reproducibility); linearity on log scale; fraction of spiked-in Hut78 cells vs PBMC; LoD; LoQ |
| SPC_00005_12_01 | SPC_00005_12 | SPC_00005 | 1 | A7 | PBMC4 | PBMC | 2000 | Hot78 cell line | 2 | 0.000999000999 | 333 | Limit of Quantitation | High complexity samples: Precision (reproducibility); linearity on log scale; fraction of spiked-in Hut78 cells vs PBMC; LoD; LoQ |
| SPC_00005_12_02 | SPC_00005_12 | SPC_00005 | 2 | C5 | PBMC4 | PBMC | 2000 | Hot78 cell line | 2 | 0.000999000999 | 333 | Limit of Quantitation | High complexity samples: Precision (reproducibility); linearity on log scale; fraction of spiked-in Hut78 cells vs PBMC; LoD; LoQ |
| SPC_00005_12_03 | SPC_00005_12 | SPC_00005 | 3 | A6 | PBMC4 | PBMC | 2000 | Hot78 cell line | 2 | 0.000999000999 | 333 | Limit of Quantitation | High complexity samples: Precision (reproducibility); linearity on log scale; fraction of spiked-in Hut78 cells vs PBMC; LoD; LoQ |
| SPC_00006_01_01 | SPC_00006_01 | SPC_00006 | 1 | A4 | PBMC5 | PBMC | 0.02 | Hot78 cell line | 2 | 0.9900990099 | 333 | Limit of Quantitation | High complexity samples: Precision (reproducibility); linearity on log scale; fraction of spiked-in Hut78 cells vs PBMC; LoD; LoQ |
| SPC_00006_01_02 | SPC_00006_01 | SPC_00006 | 2 | A7 | PBMC5 | PBMC | 0.02 | Hot78 cell line | 2 | 0.9900990099 | 333 | Limit of Quantitation | High complexity samples: Precision (reproducibility); linearity on log scale; fraction of spiked-in Hut78 cells vs PBMC; LoD; LoQ |
| SPC_00006_01_03 | SPC_00006_01 | SPC_00006 | 3 | C8 | PBMC5 | PBMC | 0.02 | Hot78 cell line | 2 | 0.9900990099 | 333 | Limit of Quantitation | High complexity samples: Precision (reproducibility); linearity on log scale; fraction of spiked-in Hut78 cells vs PBMC; LoD; LoQ |
| SPC_00006_02_01 | SPC_00006_02 | SPC_00006 | 1 | B7 | PBMC5 | PBMC | 0.2 | Hot78 cell line | 2 | 0.9090909091 | 333 | Limit of Quantitation | High complexity samples: Precision (reproducibility); linearity on log scale; fraction of spiked-in Hut78 cells vs PBMC; LoD; LoQ |
| SPC_00006_02_02 | SPC_00006_02 | SPC_00006 | 2 | C4 | PBMC5 | PBMC | 0.2 | Hot78 cell line | 2 | 0.9090909091 | 333 | Limit of Quantitation | High complexity samples: Precision (reproducibility); linearity on log scale; fraction of spiked-in Hut78 cells vs PBMC; LoD; LoQ |
| SPC_00006_02_03 | SPC_00006_02 | SPC_00006 | 3 | C6 | PBMC5 | PBMC | 0.2 | Hot78 cell line | 2 | 0.9090909091 | 333 | Limit of Quantitation | High complexity samples: Precision (reproducibility); linearity on log scale; fraction of spiked-in Hut78 cells vs PBMC; LoD; LoQ |
| SPC_00006_03_01 | SPC_00006_03 | SPC_00006 | 1 | B6 | PBMC5 | PBMC | 2 | Hot78 cell line | 2 | 0.5 | 333 | Limit of Quantitation | High complexity samples: Precision (reproducibility); linearity on log scale; fraction of spiked-in Hut78 cells vs PBMC; LoD; LoQ |
| SPC_00006_03_02 | SPC_00006_03 | SPC_00006 | 2 | B6 | PBMC5 | PBMC | 2 | Hot78 cell line | 2 | 0.5 | 333 | Limit of Quantitation | High complexity samples: Precision (reproducibility); linearity on log scale; fraction of spiked-in Hut78 cells vs PBMC; LoD; LoQ |
| SPC_00006_03_03 | SPC_00006_03 | SPC_00006 | 3 | B1 | PBMC5 | PBMC | 2 | Hot78 cell line | 2 | 0.5 | 333 | Limit of Quantitation | High complexity samples: Precision (reproducibility); linearity on log scale; fraction of spiked-in Hut78 cells vs PBMC; LoD; LoQ |
| SPC_00006_04_01 | SPC_00006_04 | SPC_00006 | 1 | A1 | PBMC5 | PBMC | 20 | Hot78 cell line | 2 | 0.09090909091 | 333 | Limit of Quantitation | High complexity samples: Precision (reproducibility); linearity on log scale; fraction of spiked-in Hut78 cells vs PBMC; LoD; LoQ |
| SPC_00006_04_02 | SPC_00006_04 | SPC_00006 | 2 | C6 | PBMC5 | PBMC | 20 | Hot78 cell line | 2 | 0.09090909091 | 333 | Limit of Quantitation | High complexity samples: Precision (reproducibility); linearity on log scale; fraction of spiked-in Hut78 cells vs PBMC; LoD; LoQ |
| SPC_00006_04_03 | SPC_00006_04 | SPC_00006 | 3 | B3 | PBMC5 | PBMC | 20 | Hot78 cell line | 2 | 0.09090909091 | 333 | Limit of Quantitation | High complexity samples: Precision (reproducibility); linearity on log scale; fraction of spiked-in Hut78 cells vs PBMC; LoD; LoQ |
| SPC_00006_05_01 | SPC_00006_05 | SPC_00006 | 1 | C2 | PBMC5 | PBMC | 200 | Hot78 cell line | 2 | 0.009900990099 | 333 | Limit of Quantitation | High complexity samples: Precision (reproducibility); linearity on log scale; fraction of spiked-in Hut78 cells vs PBMC; LoD; LoQ |
| SPC_00006_05_02 | SPC_00006_05 | SPC_00006 | 2 | A5 | PBMC5 | PBMC | 200 | Hot78 cell line | 2 | 0.009900990099 | 333 | Limit of Quantitation | High complexity samples: Precision (reproducibility); linearity on log scale; fraction of spiked-in Hut78 cells vs PBMC; LoD; LoQ |
| SPC_00006_05_03 | SPC_00006_05 | SPC_00006 | 3 | A7 | PBMC5 | PBMC | 200 | Hot78 cell line | 2 | 0.009900990099 | 333 | Limit of Quantitation | High complexity samples: Precision (reproducibility); linearity on log scale; fraction of spiked-in Hut78 cells vs PBMC; LoD; LoQ |
| SPC_00006_06_01 | SPC_00006_06 | SPC_00006 | 1 | B2 | PBMC5 | PBMC | 2000 | Hot78 cell line | 2 | 0.000999000999 | 333 | Limit of Quantitation | High complexity samples: Precision (reproducibility); linearity on log scale; fraction of spiked-in Hut78 cells vs PBMC; LoD; LoQ |
| SPC_00006_06_02 | SPC_00006_06 | SPC_00006 | 2 | B3 | PBMC5 | PBMC | 2000 | Hot78 cell line | 2 | 0.000999000999 | 333 | Limit of Quantitation | High complexity samples: Precision (reproducibility); linearity on log scale; fraction of spiked-in Hut78 cells vs PBMC; LoD; LoQ |
| SPC_00006_06_03 | SPC_00006_06 | SPC_00006 | 3 | B5 | PBMC5 | PBMC | 2000 | Hot78 cell line | 2 | 0.000999000999 | 333 | Limit of Quantitation | High complexity samples: Precision (reproducibility); linearity on log scale; fraction of spiked-in Hut78 cells vs PBMC; LoD; LoQ |
| SPC_00007_01_01 | SPC_00007_01 | SPC_00007 | 1 | C4 | B50%+T50% | PanB/PanT | 2000 | Hot78 cell line | 2 | 0.000999000999 | 333 | PanB/PanT | High complexity samples: Precision; fraction of B & T cells; fraction of V & J genes; fraction of spiked-in Hut78 cells |
| SPC_00007_01_02 | SPC_00007_01 | SPC_00007 | 3 | B8 | B50%+T50% | PanB/PanT | 2000 | Hot78 cell line | 2 | 0.000999000999 | 333 | PanB/PanT | High complexity samples: Precision; fraction of B & T cells; fraction of V & J genes; fraction of spiked-in Hut78 cells |
| SPC_00007_01_03 | SPC_00007_01 | SPC_00007 | 4 | C4 | B50%+T50% | PanB/PanT | 2000 | Hot78 cell line | 2 | 0.000999000999 | 333 | PanB/PanT | High complexity samples: Precision; fraction of B & T cells; fraction of V & J genes; fraction of spiked-in Hut78 cells |
| SPC_00007_02_01 | SPC_00007_02 | SPC_00007 | 1 | C3 | B25%+T75% | PanB/PanT | 2000 | Hot78 cell line | 2 | 0.000999000999 | 333 | PanB/PanT | High complexity samples: Precision; fraction of B & T cells; fraction of V & J genes; fraction of spiked-in Hut78 cells |
| SPC_00007_02_02 | SPC_00007_02 | SPC_00007 | 3 | C7 | B25%+T75% | PanB/PanT | 2000 | Hot78 cell line | 2 | 0.000999000999 | 333 | PanB/PanT | High complexity samples: Precision; fraction of B & T cells; fraction of V & J genes; fraction of spiked-in Hut78 cells |
| SPC_00007_02_03 | SPC_00007_02 | SPC_00007 | 4 | C7 | B25%+T75% | PanB/PanT | 2000 | Hot78 cell line | 2 | 0.000999000999 | 333 | PanB/PanT | High complexity samples: Precision; fraction of B & T cells; fraction of V & J genes; fraction of spiked-in Hut78 cells |
| SPC_00007_03_01 | SPC_00007_03 | SPC_00007 | 1 | C7 | B75%+T25% | PanB/PanT | 2000 | Hot78 cell line | 2 | 0.000999000999 | 333 | PanB/PanT | High complexity samples: Precision; fraction of B & T cells; fraction of V & J genes; fraction of spiked-in Hut78 cells |
| SPC_00007_03_02 | SPC_00007_03 | SPC_00007 | 3 | A5 | B75%+T25% | PanB/PanT | 2000 | Hot78 cell line | 2 | 0.000999000999 | 333 | PanB/PanT | High complexity samples: Precision; fraction of B & T cells; fraction of V & J genes; fraction of spiked-in Hut78 cells |
| SPC_00007_03_03 | SPC_00007_03 | SPC_00007 | 4 | A8 | B75%+T25% | PanB/PanT | 2000 | Hot78 cell line | 2 | 0.000999000999 | 333 | PanB/PanT | High complexity samples: Precision; fraction of B & T cells; fraction of V & J genes; fraction of spiked-in Hut78 cells |
| SPC_00008_01_01 | SPC_00008_01 | SPC_00008 | 1 | A8 | Pan T | Pan T | 2000 | Hot78 cell line | 2 | 0.000999000999 | 333 | PanB/PanT | High complexity samples: Precision; fraction of B & T cells; fraction of V & J genes; fraction of spiked-in Hut78 cells |
| SPC_00008_01_02 | SPC_00008_01 | SPC_00008 | 3 | A4 | Pan T | Pan T | 2000 | Hot78 cell line | 2 | 0.000999000999 | 333 | PanB/PanT | High complexity samples: Precision; fraction of B & T cells; fraction of V & J genes; fraction of spiked-in Hut78 cells |
| SPC_00008_01_03 | SPC_00008_01 | SPC_00008 | 5 | C2 | Pan T | Pan T | 2000 | Hot78 cell line | 2 | 0.000999000999 | 333 | PanB/PanT | High complexity samples: Precision; fraction of B & T cells; fraction of V & J genes; fraction of spiked-in Hut78 cells |
| SPC_00009_01_01 | SPC_00009_01 | SPC_00009 | 1 | A5 | Pan B | Pan B | 2000 | Hot78 cell line | 2 | 0.000999000999 | 333 | PanB/PanT | High complexity samples: Precision; fraction of B & T cells; fraction of V & J genes; fraction of spiked-in Hut78 cells |
| SPC_00009_01_02 | SPC_00009_01 | SPC_00009 | 3 | B6 | Pan B | Pan B | 2000 | Hot78 cell line | 2 | 0.000999000999 | 333 | PanB/PanT | High complexity samples: Precision; fraction of B & T cells; fraction of V & J genes; fraction of spiked-in Hut78 cells |
| SPC_00009_01_03 | SPC_00009_01 | SPC_00009 | 5 | B3 | Pan B | Pan B | 2000 | Hot78 cell line | 2 | 0.000999000999 | 333 | PanB/PanT | High complexity samples: Precision; fraction of B & T cells; fraction of V & J genes; fraction of spiked-in Hut78 cells |
