## Supplementary material for "immunoPETE: A DNA-based integrated B-cell and T-cell receptor profiling platform": Table S3

### Table S3 - Cell type ratio measurements in the Pan-B / Pan-T set

| | Mixture composition | | | | |
|---|---|---|---|---|---|
| | B100 | B75-T25 | B50-T50 | B25-T75 | T100 |
| **Target BCR %** | **100** | **75** | **50** | **25** | **0** |
| **Target TCR %** | **0** | **25** | **50** | **75** | **100** |
| **Detected IGH (mean)** | **99.795** | **76.647** | **51.575** | **27.324** | **0.227** |
| Detected IGH (std) | 0.028 | 0.257 | 1.402 | 3.753 | 0.009 |
| Detected IGH (CV) | 0.000 | 0.003 | 0.027 | 0.137 | 0.040 |
| **Detected TRB+TRD (mean)** | **0.205** | **23.353** | **48.425** | **72.676** | **99.773** |
| Detected TRB+TRD (std) | 0.028 | 0.257 | 1.402 | 3.753 | 0.009 |
| Detected TRB+TRD (CV) | 0.137 | 0.011 | 0.029 | 0.052 | 0.000 |
| Detected TRB (mean) | 0.195 | 22.688 | 47.101 | 70.864 | 96.849 |
| Detected TRB (std) | 0.026 | 0.227 | 1.384 | 3.279 | 0.167 |
| Detected TRB (CV) | 0.133 | 0.010 | 0.029 | 0.046 | 0.002 |
| Detected TRD (mean) | 0.01 | 0.665 | 1.324 | 1.813 | 2.924 |
| Detected TRD (std) | 0.004 | 0.03 | 0.025 | 0.476 | 0.162 |
| Detected TRD (CV) | 0.400 | 0.045 | 0.019 | 0.263 | 0.055 |
