## Supplementary material for "immunoPETE: A DNA-based integrated B-cell and T-cell receptor profiling platform": Table S4

### Table S4 - Sample Summary Statistics of PBMC1 and PBMC2

| | PBMC1_mean | PBMC1_std | PBMC1_cv | PBMC2_mean | PBMC2_std | PBMC2_cv |
|---|---|---|---|---|---|---|
| **valid_on_target_reads** | 2569543.875 | 797768.346 | 0.31 | 3057235.375 | 200328.193 | 0.066 |
| **UMI_families** | 90534.625 | 14278.24 | 0.158 | 105337.125 | 5935.124 | 0.056 |
| **percent_functional** | 80.391 | 0.563 | 0.007 | 81.942 | 0.223 | 0.003 |
| **total_cell_count** | 72716.125 | 11144.182 | 0.153 | 86306.75 | 4700.44 | 0.054 |
| **TRB** | 62772.375 | 9764.853 | 0.156 | 70760.5 | 3879.762 | 0.055 |
| **TRD** | 3082.875 | 782.457 | 0.254 | 1535.625 | 217.478 | 0.142 |
| **IGH** | 6860.875 | 640.001 | 0.093 | 14010.625 | 666.304 | 0.048 |
| **TRB_percent** | 86.289 | 0.387 | 0.004 | 81.986 | 0.278 | 0.003 |
| **TRD_percent** | 4.162 | 0.62 | 0.149 | 1.772 | 0.158 | 0.089 |
| **IGH_percent** | 9.551 | 0.895 | 0.094 | 16.242 | 0.338 | 0.021 |
| **clone_count** | 59154.125 | 8474.219 | 0.143 | 74638.125 | 3671.829 | 0.049 |
| **D50** | 0.386 | 0.009 | 0.024 | 0.422 | 0.003 | 0.008 |
| **gini_index** | 0.181 | 0.012 | 0.066 | 0.132 | 0.005 | 0.035 |
