## Supplementary material for "immunoPETE: A DNA-based integrated B-cell and T-cell receptor profiling platform": Table S5

### Table S5 - Identity and frequency of top 10 clones in PBMC1-4

| | Summary statistics | | | Individual replicates | | | | | | | | | | | |
|---|---|---|---|---|---|---|---|---|---|---|---|---|---|---|---|
| **PBMC1** | **mean** | **std** | **cv** | SPC_00002_01_01_sg86 | SPC_00002_01_02_sg99 | SPC_00002_02_01_sg74 | SPC_00002_02_02_sg112 | SPC_00002_03_01_sg81 | SPC_00002_03_02_sg118 | SPC_00002_04_01_sg79 | SPC_00002_04_02_sg98 | | | | |
| TRBV3-1__CASSQDRFSGNTIYF__TRBJ1-3 | 0.012349 | 0.000371 | 0.030073 | 0.011914 | 0.012309 | 0.012822 | 0.011993 | 0.011942 | 0.012505 | 0.012502 | 0.012808 | | | | |
| TRBV7-9__CASSSTGRERDEQFF__TRBJ2-1 | 0.005065 | 0.001238 | 0.244385 | 0.006011 | 0.005961 | 0.005641 | 0.004345 | 0.005762 | 0.002335 | 0.005535 | 0.004928 | | | | |
| TRBV4-1__CASSVGREASPLHF__TRBJ1-6 | 0.004424 | 0.000943 | 0.213129 | 0.004307 | 0.004161 | 0.003976 | 0.003931 | 0.004172 | 0.006722 | 0.004270 | 0.003851 | | | | |
| TRBV20-1__CSAELAGGFEQYF__TRBJ2-7 | 0.003169 | 0.000288 | 0.090866 | 0.003582 | 0.003387 | 0.003013 | 0.003129 | 0.003390 | 0.002641 | 0.003119 | 0.003091 | | | | |
| TRAV29DV5__CAASTLIFPRGLYTDKLIF__TRDJ1 | 0.002627 | 0.000721 | 0.274468 | 0.002402 | 0.002974 | 0.002697 | 0.002875 | 0.002907 | 0.000938 | 0.003145 | 0.003078 | | | | |
| TRBV12-3__CASSFFRGGSSYEQYF__TRBJ2-7 | 0.002467 | 0.000354 | 0.143588 | 0.002603 | 0.002574 | 0.002476 | 0.002313 | 0.002751 | 0.001659 | 0.002659 | 0.002698 | | | | |
| TRBV4-1__CASSQEDSVVAYEQYF__TRBJ2-7 | 0.002451 | 0.000208 | 0.085051 | 0.002563 | 0.002599 | 0.002408 | 0.002647 | 0.002321 | 0.002008 | 0.002569 | 0.002496 | | | | |
| TRBV7-9__CASSYTGSANTGELFF__TRBJ2-2 | 0.002379 | 0.000177 | 0.074227 | 0.002455 | 0.002487 | 0.002284 | 0.002580 | 0.002021 | 0.002488 | 0.002288 | 0.002432 | | | | |
| TRBV6-4__CASSDSTGANVLTF__TRBJ2-6 | 0.002330 | 0.000310 | 0.132973 | 0.001999 | 0.002012 | 0.002710 | 0.002447 | 0.002386 | 0.002597 | 0.001917 | 0.002572 | | | | |
| TRBV6-4__CASSDGTSGSGEQFF__TRBJ2-1 | 0.001891 | 0.000453 | 0.239335 | 0.002254 | 0.001999 | 0.002036 | 0.001658 | 0.001969 | 0.000960 | 0.002467 | 0.001786 | | | | |
| **PBMC2** | **mean** | **std** | **cv** | SPC_00003_01_01_sg87 | SPC_00003_01_02_sg109 | SPC_00003_02_01_sg84 | SPC_00003_02_02_sg116 | SPC_00003_03_01_sg78 | SPC_00003_03_02_sg113 | SPC_00003_04_01_sg77 | SPC_00003_04_02_sg101 | | | | |
| TRBV18__CASSPQDPGGNQPQHF__TRBJ1-5 | 0.003377 | 0.000243 | 0.071969 | 0.003123 | 0.003106 | 0.003705 | 0.003762 | 0.003269 | 0.003373 | 0.003291 | 0.003385 | | | | |
| TRBV5-1__CASSYGQADLHF__TRBJ1-6 | 0.001314 | 0.000111 | 0.084333 | 0.001452 | 0.001465 | 0.001318 | 0.001279 | 0.001150 | 0.001300 | 0.001357 | 0.001195 | | | | |
| TRBV9__CASTSPGGMAFF__TRBJ1-1 | 0.001299 | 0.000171 | 0.131412 | 0.001175 | 0.001187 | 0.001473 | 0.001117 | 0.001353 | 0.001107 | 0.001504 | 0.001475 | | | | |
| TRBV20-1__CSARDRVTEAFF__TRBJ1-1 | 0.001210 | 0.000138 | 0.113890 | 0.000991 | 0.001035 | 0.001389 | 0.001217 | 0.001308 | 0.001321 | 0.001199 | 0.001218 | | | | |
| TRBV12-3__CASSPAGLGGTEAFF__TRBJ1-1 | 0.001152 | 0.000229 | 0.199035 | 0.001291 | 0.000795 | 0.001389 | 0.000844 | 0.001308 | 0.001203 | 0.001335 | 0.001050 | | | | |
| TRBV20-1__CSAPQVRGSWGYTF__TRBJ1-2 | 0.001028 | 0.000091 | 0.088361 | 0.001118 | 0.001086 | 0.001009 | 0.001043 | 0.000845 | 0.000956 | 0.001063 | 0.001106 | | | | |
| TRBV6-4__CATRDRDNSPLHF__TRBJ1-6 | 0.000955 | 0.000110 | 0.115470 | 0.001037 | 0.000770 | 0.000974 | 0.000832 | 0.000936 | 0.000988 | 0.000984 | 0.001117 | | | | |
| TRBV4-3__CASSQSVVQGGITGELFF__TRBJ2-2 | 0.000960 | 0.000089 | 0.092312 | 0.001049 | 0.001086 | 0.000867 | 0.000906 | 0.000868 | 0.000892 | 0.001041 | 0.000972 | | | | |
| TRBV5-4__CASSLAGGSLRNEQFF__TRBJ2-1 | 0.000954 | 0.000140 | 0.146622 | 0.000922 | 0.000909 | 0.000879 | 0.001180 | 0.000879 | 0.000999 | 0.000746 | 0.001117 | | | | |
| TRBV19__CASSTGDSNQPQHF__TRBJ1-5 | 0.000877 | 0.000125 | 0.141963 | 0.000910 | 0.000997 | 0.000808 | 0.000670 | 0.000823 | 0.000988 | 0.001029 | 0.000793 | | | | |
| **PBMC3** | **mean** | **std** | **cv** | SPC_00004_01_01_sg28 | SPC_00004_02_01_sg43 | SPC_00004_02_02_sg68 | SPC_00004_03_01_sg41 | SPC_00004_03_02_sg63 | SPC_00004_04_02_sg69 | SPC_00004_05_01_sg27 | SPC_00004_05_02_sg51 | SPC_00004_06_02_sg50 | | | |
| TRBV18__CASSPQDPGGNQPQHF__TRBJ1-5 | 0.003201 | 0.000343 | 0.107152 | 0.002653 | 0.003206 | 0.003461 | 0.003203 | 0.003759 | 0.002982 | 0.002832 | 0.003476 | 0.003239 | | | |
| TRBV20-1__CSARDRVTEAFF__TRBJ1-1 | 0.001339 | 0.000184 | 0.137154 | 0.001222 | 0.001702 | 0.001199 | 0.001248 | 0.001117 | 0.001252 | 0.001522 | 0.001392 | 0.001398 | | | |
| TRBV5-1__CASSYGQADLHF__TRBJ1-6 | 0.001239 | 0.000161 | 0.129599 | 0.000873 | 0.001258 | 0.001363 | 0.001331 | 0.001439 | 0.001208 | 0.001181 | 0.001210 | 0.001291 | | | |
| TRBV12-3__CASSPAGLGGTEAFF__TRBJ1-1 | 0.001239 | 0.000252 | 0.202995 | 0.001222 | 0.001726 | 0.001090 | 0.001560 | 0.000988 | 0.000984 | 0.001273 | 0.001143 | 0.001169 | | | |
| TRBV9__CASTSPGGMAFF__TRBJ1-1 | 0.001166 | 0.000153 | 0.131637 | 0.000838 | 0.001282 | 0.001117 | 0.001248 | 0.001074 | 0.001088 | 0.001328 | 0.001248 | 0.001268 | | | |
| TRBV20-1__CSAPQVRGSWGYTF__TRBJ1-2 | 0.000975 | 0.000154 | 0.157658 | 0.001082 | 0.000863 | 0.001226 | 0.000770 | 0.000795 | 0.001118 | 0.000922 | 0.001037 | 0.000962 | | | |
| TRBV5-4__CASSLAGGSLRNEQFF__TRBJ2-1 | 0.000968 | 0.000153 | 0.158034 | 0.000838 | 0.000888 | 0.001308 | 0.000998 | 0.001010 | 0.001058 | 0.000876 | 0.000931 | 0.000802 | | | |
| TRBV6-4__CATRDRDNSPLHF__TRBJ1-6 | 0.000852 | 0.000134 | 0.157349 | 0.000908 | 0.000814 | 0.000818 | 0.000624 | 0.000795 | 0.000954 | 0.001033 | 0.000720 | 0.001001 | | | |
| TRBV19__CASSTGDSNQPQHF__TRBJ1-5 | 0.000824 | 0.000131 | 0.159293 | 0.000558 | 0.000765 | 0.000790 | 0.000894 | 0.000709 | 0.000969 | 0.000885 | 0.000903 | 0.000940 | | | |
| TRBV4-3__CASSQSVVQGGITGELFF__TRBJ2-2 | 0.000865 | 0.000099 | 0.114728 | 0.000698 | 0.000839 | 0.000845 | 0.000977 | 0.000967 | 0.000939 | 0.000756 | 0.000951 | 0.000817 | | | |
| **PBMC4** | **mean** | **std** | **cv** | SPC_00005_01_01_sg82 | SPC_00005_01_02_sg120 | SPC_00005_02_01_sg91 | SPC_00005_02_02_sg102 | SPC_00005_03_01_sg88 | SPC_00005_03_02_sg97 | SPC_00005_04_01_sg76 | SPC_00005_04_02_sg111 | SPC_00005_05_01_sg94 | SPC_00005_05_02_sg100 | SPC_00005_06_01_sg90 | SPC_00005_06_02_sg115 |
| TRBV20-1__CSARQVAGDTSPLHF__TRBJ1-6 | 0.009297 | 0.000535 | 0.057601 | 0.008957 | 0.010609 | 0.008751 | 0.009958 | 0.009195 | 0.009111 | 0.009302 | 0.009612 | 0.008905 | 0.009287 | 0.009050 | 0.008823 |
| TRBV14__CASSQVGSSFQWGYTF__TRBJ1-2 | 0.007476 | 0.000533 | 0.071246 | 0.007077 | 0.008316 | 0.007869 | 0.008210 | 0.006662 | 0.007995 | 0.007322 | 0.007666 | 0.007311 | 0.007202 | 0.007249 | 0.006830 |
| TRBV30__CAWSVQIAGPYEQYF__TRBJ2-7 | 0.004091 | 0.000470 | 0.114903 | 0.003829 | 0.002798 | 0.003830 | 0.004393 | 0.004057 | 0.004131 | 0.004522 | 0.004111 | 0.004361 | 0.004245 | 0.004557 | 0.004260 |
| TRBV6-5__CALGTGPWGEQYF__TRBJ2-7 | 0.003567 | 0.000272 | 0.076330 | 0.003043 | 0.003769 | 0.003760 | 0.003956 | 0.003455 | 0.003714 | 0.003327 | 0.003736 | 0.003417 | 0.003672 | 0.003211 | 0.003741 |
| TRBV6-2__CASSLAGEAYNEQFF__TRBJ2-1 | 0.003131 | 0.000366 | 0.116802 | 0.003795 | 0.002992 | 0.003412 | 0.002990 | 0.003384 | 0.003448 | 0.002695 | 0.002841 | 0.003261 | 0.003315 | 0.002517 | 0.002919 |
| TRBV12-3__CASSTSSGGLSGYTF__TRBJ1-2 | 0.003009 | 0.000206 | 0.068288 | 0.003487 | 0.003109 | 0.003018 | 0.003220 | 0.002782 | 0.002698 | 0.002882 | 0.002914 | 0.003060 | 0.002982 | 0.003021 | 0.002941 |
| TRBV6-2__CASSYPTGGNSPLHF__TRBJ1-6 | 0.002872 | 0.000297 | 0.103525 | 0.002769 | 0.003070 | 0.003644 | 0.002806 | 0.002693 | 0.002615 | 0.002484 | 0.003035 | 0.002913 | 0.002733 | 0.002741 | 0.002963 |
| TRBV19__CASSTTGGTGGEKLFF__TRBJ1-4 | 0.002569 | 0.000338 | 0.131403 | 0.002940 | 0.002681 | 0.002994 | 0.002415 | 0.002799 | 0.002865 | 0.002777 | 0.001947 | 0.002336 | 0.002384 | 0.002608 | 0.002082 |
| TRBV4-3__CASSQATGRYEQYF__TRBJ2-7 | 0.002218 | 0.000211 | 0.095011 | 0.002085 | 0.002448 | 0.001834 | 0.002277 | 0.002551 | 0.002298 | 0.001910 | 0.002442 | 0.002199 | 0.002226 | 0.002180 | 0.002171 |
| TRBV6-2__CASSYPTSGFSEQFF__TRBJ2-1 | 0.001844 | 0.000241 | 0.130941 | 0.002188 | 0.001865 | 0.001834 | 0.001495 | 0.002091 | 0.002165 | 0.001804 | 0.001656 | 0.001860 | 0.002010 | 0.001451 | 0.001711 |
