## Supplementary material for "immunoPETE: A DNA-based integrated B-cell and T-cell receptor profiling platform": Table S6

### Table S6 - Chi-squared test results for V and J gene pairing independence

| | TRB_chi2 | TRB_p_value | TRB_cramer_v | TRD_chi2 | TRD_p_value | TRD_cramer_v | IGH_chi2 | IGH_p_value | IGH_cramer_v |
|---|---|---|---|---|---|---|---|---|---|
| **PBMC1** | 35411.02374 | 0 | 0.1483691433 | 7.75E+02 | 2.07E-150 | 0.2007622579 | 765.795145 | 1.12E-65 | 0.1042225622 |
| **PBMC2** | 9781.079881 | 0 | 0.07749142668 | 2.08E+02 | 1.03E-32 | 0.1584607339 | 1720.575747 | 2.11E-237 | 0.1118709012 |
| **PBMC3** | 12613.20095 | 0 | 0.07765458657 | 249.4849163 | 5.21E-41 | 0.1472579788 | 2166.626534 | 0 | 0.1122136682 |
| **PBMC4** | 31690.31173 | 0 | 0.1214398482 | 291.6761261 | 1.57E-49 | 0.1541047038 | 2249.668528 | 0 | 0.09850335742 |
| **PBMC5** | 22755.17115 | 0 | 0.1287636769 | 2.53E+02 | 9.04E-42 | 0.1615708455 | 1413.138836 | 2.63E-179 | 0.09920771377 |
