## Supplementary material for "immunoPETE: A DNA-based integrated B-cell and T-cell receptor profiling platform": Table S7

| **Table S7 - Demographic and clinical description of the NMIBC cohort in the case study** | | |
|---|---|---|
| Characteristic | Median (range) or Number of patients (%) | |
| Median age | 66.68 (39.41 to 79.89) | |
| Median follow up duration | 1890 days (292 to 2949) | |
| Sex | | |
| | Male | 22 (91.6%) |
| | Female | 2 (8.3%) |
| Stage at diagnosis | | |
| | Ta | 3 (12.5%) |
| | T1 | 21 (87.5%) |
| Grade at diagnosis | | |
| | G2 | 7 (29.2%) |
| | G3 | 17 (70.8%) |
| Carcinoma *in situ* present | | |
| | Yes | 8 (33.3%) |
| | No | 11 (45.8%) |
| | Unknown | 5 (20.8%) |
| EAU Risk category | | |
| | Intermediate | 1 (4.1%) |
| | High | 23 (95.8%) |
| Treatment in addition to TURBT | | |
| | BCG | 22 (91.6%) |
| | None | 1 (4.1%) |
| | Unknown | 1 (4.1%) |
| Outcomes | | |
| | Recurrence only | 7 (29.2%) |
| | Recurrence and progression | 8 (33.3%) |
| | No recurrence/progression | 9 (37.5%) |
