## Supplementary material for "immunoPETE: A DNA-based integrated B-cell and T-cell receptor profiling platform": Table S8

### Table S8 - Hill Numbers for PBMC1-4

| | | | Order | | | | | | |
|---|---|---|---|---|---|---|---|---|---|
| **sample_name** | **dna_source** | **dna_source_ng** | **0** | **1** | **2** | **3** | **4** | **5** | **inf** |
| SPC_00002_01_01_sg86 | PBMC1 | 2000 | 60799 | 40760.76174 | 3319.317841 | 679.978095 | 355.6671961 | 251.3540718 | 83.93693694 |
| SPC_00002_01_02_sg99 | PBMC1 | 2000 | 64789 | 42652.96678 | 3219.156564 | 655.351451 | 342.1885516 | 241.8004116 | 81.24060914 |
| SPC_00002_02_01_sg74 | PBMC1 | 2000 | 58811 | 38881.16151 | 3140.229854 | 630.6659451 | 326.7669739 | 230.5082214 | 77.98927039 |
| SPC_00002_02_02_sg112 | PBMC1 | 2000 | 60887 | 41080.13459 | 3553.447363 | 701.451297 | 358.6942934 | 251.090751 | 83.38238573 |
| SPC_00002_03_01_sg81 | PBMC1 | 2000 | 62046 | 41217.95463 | 3347.712118 | 683.5155276 | 356.008491 | 251.0812598 | 83.73799127 |
| SPC_00002_03_02_sg118 | PBMC1 | 2000 | 38727 | 28526.60316 | 3386.082313 | 649.3641789 | 334.487425 | 236.4284055 | 79.96684119 |
| SPC_00002_04_01_sg79 | PBMC1 | 2000 | 63289 | 41519.38052 | 3176.614092 | 647.3555026 | 336.9254323 | 237.7090804 | 79.98773006 |
| SPC_00002_04_02_sg98 | PBMC1 | 2000 | 63885 | 42121.12048 | 3215.871582 | 637.9763143 | 328.7526519 | 231.2835395 | 78.07912957 |
| SPC_00003_01_01_sg87 | PBMC2 | 2000 | 75244 | 61831.18747 | 19930.71077 | 4444.245281 | 2060.48399 | 1331.631311 | 320.2398524 |
| SPC_00003_01_02_sg109 | PBMC2 | 2000 | 69083 | 57244.37045 | 19706.27507 | 4519.078852 | 2086.15121 | 1343.412626 | 321.9796748 |
| SPC_00003_02_01_sg84 | PBMC2 | 2000 | 73090 | 60066.60378 | 18133.43672 | 3706.556788 | 1679.79208 | 1083.645839 | 269.8878205 |
| SPC_00003_02_02_sg116 | PBMC2 | 2000 | 69941 | 57691.49196 | 18237.64212 | 3729.020801 | 1664.975725 | 1067.526109 | 265.8085809 |
| SPC_00003_03_01_sg78 | PBMC2 | 2000 | 76572 | 62600.18316 | 19796.88096 | 4303.010175 | 1965.167513 | 1263.68825 | 305.9241379 |
| SPC_00003_03_02_sg113 | PBMC2 | 2000 | 79714 | 64420.29372 | 18896.94275 | 4100.490473 | 1888.072995 | 1216.579838 | 296.4299363 |
| SPC_00003_04_01_sg77 | PBMC2 | 2000 | 76267 | 61926.6863 | 18672.11637 | 4127.839537 | 1923.892716 | 1247.448514 | 303.8316151 |
| SPC_00003_04_02_sg101 | PBMC2 | 2000 | 77194 | 62709.08712 | 18843.05817 | 4073.737582 | 1874.572119 | 1209.640434 | 295.3960396 |
| SPC_00004_01_01_sg28 | PBMC3 | 500 | 25879 | 23249.99251 | 14315.25597 | 5099.553941 | 2486.099438 | 1617.626497 | 376.9605263 |
| SPC_00004_02_01_sg43 | PBMC3 | 750 | 36302 | 31832.44189 | 15520.64836 | 4156.781874 | 1951.459595 | 1273.987305 | 311.8923077 |
| SPC_00004_02_02_sg68 | PBMC3 | 750 | 33114 | 29262.15996 | 14837.53825 | 3989.42546 | 1830.020789 | 1178.858737 | 288.9527559 |
| SPC_00004_03_01_sg41 | PBMC3 | 1000 | 42684 | 36773.12321 | 16219.49088 | 4212.76986 | 1981.555341 | 1287.35493 | 312.2272727 |
| SPC_00004_03_02_sg63 | PBMC3 | 1000 | 41513 | 35941.64515 | 15909.87444 | 3748.630862 | 1669.529793 | 1068.894485 | 266.0228571 |
| SPC_00004_04_02_sg69 | PBMC3 | 1500 | 58997 | 49817.7905 | 19385.36542 | 4710.715633 | 2190.291419 | 1411.884707 | 335.385 |
| SPC_00004_05_01_sg27 | PBMC3 | 2000 | 93078 | 75135.5241 | 21571.30458 | 4796.344743 | 2276.136873 | 1484.989589 | 353.1530945 |
| SPC_00004_05_02_sg51 | PBMC3 | 2000 | 89669 | 72396.40392 | 19817.90179 | 4018.634134 | 1824.622871 | 1173.46383 | 287.6657459 |
| SPC_00004_06_02_sg50 | PBMC3 | 2500 | 111037 | 87269.60225 | 21200.66351 | 4315.213436 | 1981.203104 | 1276.630892 | 308.7570755 |
| SPC_00005_01_01_sg82 | PBMC4 | 500 | 24181 | 17960.95973 | 3473.91405 | 854.5671379 | 466.6485596 | 336.0951619 | 111.6412214 |
| SPC_00005_01_02_sg120 | PBMC4 | 500 | 21444 | 15942.00533 | 2958.800626 | 702.5243966 | 380.5420859 | 274.4615311 | 94.26373626 |
| SPC_00005_02_01_sg91 | PBMC4 | 750 | 34888 | 24780.55483 | 3530.583267 | 827.4654146 | 454.3924426 | 329.5580693 | 114.270557 |
| SPC_00005_02_02_sg102 | PBMC4 | 750 | 35356 | 25037.06652 | 3257.026471 | 737.5892054 | 402.2949108 | 291.3321611 | 100.4203233 |
| SPC_00005_03_01_sg88 | PBMC4 | 1000 | 45016 | 31038.71898 | 3797.858189 | 861.531363 | 464.984041 | 332.7212824 | 108.7495183 |
| SPC_00005_03_02_sg97 | PBMC4 | 1000 | 47427 | 32024.48805 | 3544.20282 | 803.0586716 | 438.5220078 | 317.4629214 | 109.76234 |
| SPC_00005_04_01_sg76 | PBMC4 | 1500 | 66165 | 42886.21555 | 3812.26278 | 829.0693871 | 447.1483558 | 321.4226771 | 107.5025189 |
| SPC_00005_04_02_sg111 | PBMC4 | 1500 | 64508 | 42101.33155 | 3664.140528 | 791.7863255 | 427.6424014 | 307.908926 | 104.0352201 |
| SPC_00005_05_01_sg94 | PBMC4 | 2000 | 82809 | 51315.45779 | 3874.422587 | 853.0891385 | 463.8053816 | 334.4167706 | 112.3024691 |
| SPC_00005_05_02_sg100 | PBMC4 | 2000 | 90604 | 55653.18201 | 3853.397966 | 831.4033271 | 449.6679987 | 323.3110564 | 107.6717352 |
| SPC_00005_06_01_sg90 | PBMC4 | 2500 | 106122 | 63675.35227 | 4004.929076 | 851.9033716 | 459.6331555 | 330.5454935 | 110.4941905 |
| SPC_00005_06_02_sg115 | PBMC4 | 2500 | 101812 | 62149.2127 | 4151.777144 | 888.9221708 | 479.5332091 | 344.1566084 | 113.3358522 |
